# Lysosomes cell autonomously regulate myeloid cell states and immune responses

**DOI:** 10.1101/2024.11.11.623074

**Authors:** Leon Tejwani, Christopher Balak, Lukas L. Skuja, David Tatarakis, Anil Rana, Gabriel A. Fitzgerald, Connie Ha, Gabrielly Lunkes de Melo, Elizabeth W. Sun, Catherine M. Heffner, Matthew J. Simon, Jason C. Dugas, Lily Sarrafha, Eric Liang, Thomas Sandmann, Christopher K. Glass, Gilbert Di Paolo

**Affiliations:** Denali Therapeutics Inc., South San Francisco, CA, USA; Department of Medicine, University of California San Diego, La Jolla, CA, USA; Department of Cellular and Molecular Medicine, University of California San Diego, La Jolla, CA, USA

**Keywords:** progranulin, lysosomes, microglia, macrophages, frontotemporal dementia, frontotemporal lobar degeneration, lysosomal pH, single-cell RNA-seq, neurodegeneration, disease-associated microglia, epigenetics, enhancers

## Abstract

Myeloid cells maintain tissue homeostasis via the recognition, engulfment, and lysosomal clearance of dying cells and cellular debris, which is often accompanied by changes from homeostatic to reactive states. While a role for phagocytic receptors in gating these transitions has been described^1,2^, less is known about if and how lysosomes can contribute to transcriptional and functional plasticity. To determine how lysosomal health impacts myeloid cell states, we evaluated microglia and macrophages deficient for progranulin (encoded by *Grn*), a lysosomal protein with pleiotropic functions whose loss is associated with several neurodegenerative diseases^3–8^. Single-cell RNA-sequencing of the aged mouse brain identified a *Grn* knockout (KO)-specific microglial subpopulation marked by high GPNMB expression that displays hallmarks of lysosomal dysfunction, including lipofuscin accumulation. Epigenetic analysis of aged microglia revealed MITF/TFE transcription factors as key mediators of the transcriptional states associated with *Grn* deficiency. In addition to identifying a core myeloid cell transcriptional response to diverse lysosomal stressors, targeted perturbations of various lysosomal properties *in vitro* uncovered a cell autonomous, TREM2- independent, response to lysosomal deacidification (via v-ATPase or VPS34 loss of function) that overlaps with *Grn* KO microglia phenotypes, including the induction of a lysosomal gene program, increased proliferation, and secretion of pro-inflammatory cytokines. Compound loss-of-function approaches established GPNMB upregulation upon lysosomal stress is required for the compensatory response to enhance lysosomal function via promoting acidification. Finally, pharmacological endolysosomal reacidification through sodium/proton exchanger inhibition partially rescued *Grn* KO microglia phenotypes. Overall, these data establish a fundamental link between lysosomal health and myeloid cell epigenetic, transcriptional, and functional states observed in neurodegeneration models.

## MAIN

The ability of macrophages and microglia to appropriately function in their respective tissues is contingent on their capacity to sense extracellular cues and mount responses that involve chemotaxis, proliferation, cytokine and chemokine secretion, and phagocytosis of various cargos. These alterations are often associated with a remodeling of transcriptional, epigenetic, metabolomic, proteomic, and morphological profiles, which collectively contribute to diverse microglial states^9^. In aging and in neurodegenerative diseases, microglia are presented with numerous extracellular challenges, including dead neurons, myelin and synaptic debris, and aberrant protein aggregates^1^. As the primary phagocytes of the CNS, microglia play an essential role in the clearance of these neurodegeneration-associated molecular patterns (NAMPs), which requires recognition via a battery of cell surface receptors, engulfment through phagocytosis, and subsequent lysosomal degradation^1^. Several phagocytic receptors that recognize NAMPs, such as TREM2, have been shown to be obligatory for the functional and transcriptional conversion of homeostatic microglia to specific disease-associated microglial (DAM) states in certain contexts, particularly those observed in Alzheimer’s Disease (AD)^10–12^. Mechanistically, although several intracellular signaling molecules proximal to phagocytic receptors have been shown to transduce these extracellular cues^13,14^, how downstream processes along the phagolysosomal axis can influence myeloid cell states remains to be determined.

In many genetic lysosomal storage disorders (LSDs), microglia and peripheral macrophages display reactive profiles, which, in some cases, can precede overt cell loss^15,16^. Whether these states arise solely as a response to parenchymal damage or involve cell autonomous myeloid cell responses to impaired lysosomal function has not been comprehensively addressed. Notably, many neurodegenerative disease-causing genes encode proteins with primary roles related to endolysosomal function and/or the autophagy-lysosome pathway^17,18^, including *GRN* (encoding progranulin)^3–5,8,19^. In humans, haploinsufficiency of *GRN* causes frontotemporal dementia (FTD)^3,4^ and homozygous loss of function mutations cause neuronal ceroid lipofuscinosis (NCL)^5^, a LSD characterized by severe neurodegeneration. Additionally, common single nucleotide polymorphisms (SNPs) in *GRN* have been associated with AD^20,21^ and Parkinson’s

Disease (PD)^22^. *Grn* is mainly expressed by neurons and microglia in the CNS^23^ and is highly upregulated in microglia upon activation during CNS injury^24,25^ and neurodegeneration^11,26^. Although progranulin can be secreted and modulate inflammation^27–30^, neuronal outgrowth and neuroprotection^31,32^, its essential roles in regulating many aspects of lysosomal biology have come into the spotlight in recent years^8,19,33^. Therefore, given the indispensable role of the lysosome in the terminal degradation of intracellular material and extracellular cargo and the particularly high lysosomal demand in microglia and macrophages, here, we interrogated the interplay between lysosomes and inflammation. In this study, we demonstrate that persistent disruption of lysosomal homeostasis in microglia and macrophages is sufficient to trigger chronic, nonresolving sterile inflammation reminiscent to that observed during neurodegeneration through cell autonomous epigenetic remodeling of gene expression.

### scRNA-seq of aged *Grn* KO microglia

Previous bulk and single-nucleus RNA-sequencing (snRNA-seq) analyses of *Grn* KO mouse brains have identified robust age-dependent alterations in overall microglial gene expression^26,34,35^; however, due to the limited number of aged microglia previously sampled by snRNA-seq, whether these changes reflect global changes in microglia or increased microglial heterogeneity remains to be determined. To interrogate the transcriptional diversity of progranulin-deficient microglia *in vivo*, we purified CD11b^+^/CD45^+^/CD44^-^/CD163^-^/Ly6G^-^ cells from aged (24-month) WT and *Grn* KO mice and performed droplet-based scRNA-seq (**Fig. 1a,b and Extended Data** Fig. 1a). *P2ry12*- and *Tmem119*-expressing microglia (26,605 cells) were subclustered for downstream analyses, yielding ten subpopulations (MG1-MG10) (**Fig. 1c, Extended Data** Fig. 1b-f**, Supplementary Table 1**), including three homeostatic microglia clusters (MG1, MG2, and MG3; *P2ry12*^high^/*Tmem119*^high^), an undefined microglia cluster (MG4; *P2ry12*^high^/*Tmem119*^low^/*Nav2*^high^), a ribosomal microglia cluster (MG5; *Rlp35*^high^/*Rpl36a*^high^), an intermediate DAM-like cluster (MG6; *Axl*^high^/*Lpl*^low^/*Gpnmb*^low^), a canonical DAM-like cluster (MG7; *Axl*^high^/*Lpl*^high^/*Ccl4*^high^/*Gpnmb*^low^), a *Grn* KO-specific cluster (MG8; *Gpnmb*^high^/*Ms4a7*^high^/*Lpl*^low^), an interferon-responsive cluster (MG9; *Ifitm3*^high^/*Isg15*^high^), and a proliferative population (MG10; *Top2a*^high^/*Mki67*^high^). As expected with a multiplexed scRNA-seq approach to minimize batch effects, microglia from all ten mice were well-integrated in low-dimensional embeddings (**Extended Data** Fig. 1g).

**Figure 1.**
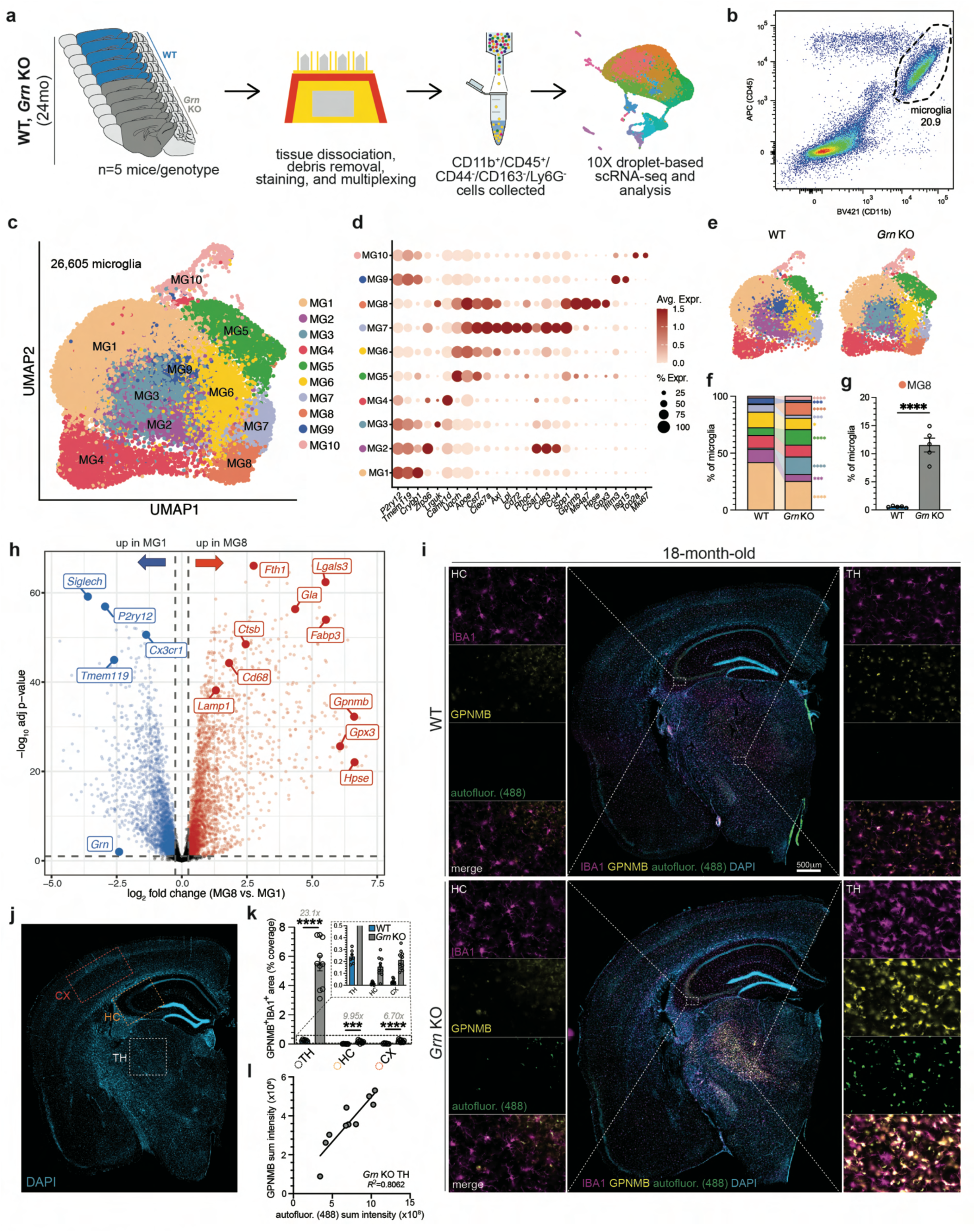
Identification of a *Grn* KO-specific microglia population. **(a)** Schematic outlining approach to enrich for CD45^+^/CD11b^+^/CD44^-^/CD163^-^/Ly6G^-^ microglia from aged (24-month) WT and *Grn* KO mouse brains for scRNA-seq (n=5 mice/genotype). **(b)** Representative FACS plot displaying CD45^+^/CD11b^+^ cell population collected for scRNA- seq. **(c)** UMAP plot of microglial subclusters identified among all high-quality microglia (26,605 cells). **(d)** Dot plot of marker genes for different microglia subclusters. **(e)** UMAP of microglial subclusters, separated by genotype (9,000 microglia/genotype). **(f)** Relative percent abundance of microglial subclusters across genotypes. **(g)** Bar graph displaying percentage of MG8 microglia in each genotype (n=5 mice/genotype). **(h)** Volcano plot of differential expression between MG8 and homeostatic MG1 population identified in *Grn* KO mice. **(i)** Representative images of coronal sections from 18-month WT and *Grn* KO mice stained for DAPI, IBA1 (microglia), and GPNMB (MG8 marker). Insets from the hippocampus (HC) and thalamus (TH) are displayed. **(j)** Representative image of coronal section from an 18-month WT mouse stained for DAPI to display quantified regions for cortex (CX), HC, and TH. **(k)** Quantification of % coverage of total area occupied by GPNMB^+^/IBA1^+^ microglia (WT, n=7 mice/genotype; *Grn* KO, n=10 mice/genotype). **(l)** Correlation between autofluorescence (488) and GPNMB sum intensity within the *Grn* KO TH (n=10 mice). Data presented in (g,k) are mean±s.e.m. The *propeller* R package was used to compare the relative abundance of each cluster across genotypes in (f). Student’s t-tests were performed to compare across genotypes in (g,k). \**P*<0.05, \*\**P*<0.01, \*\*\**P*<0.001, \*\*\*\**P*<0.0001.

Analysis of relative subcluster abundance revealed alterations in proportions of nearly all microglial subpopulations between genotypes (**Fig. 1e-g and Extended Data** Fig. 1h,i). Notably, MG8 was almost completely limited to *Grn* KO animals, comprising approximately 10% of all microglia compared to less than 0.5% in WT age-matched controls (**Fig. 1f,g and Extended Data** Fig. 1h-i). Differential expression analysis between MG8 and MG1 (homeostatic MG) identified several candidate marker genes for MG8, including *Gpnmb*, *Hpse*, *Gpx3*, *Fabp3*, among others (**Fig. 1h, Extended Data** Fig. 1j**, and Supplementary Table 1**). Using a cohort of 18-month WT and *Grn* KO mice, we validated the specificity of *Gpnmb*^high^ microglia for *Grn* KO mice (**Fig. 1i-k**), and determined by immunohistochemistry that the vast majority of MG8 is localized to the thalamus, a region that is preferentially impacted in *Grn* KO mice and preclinical progranulin mutation carriers^34,36,37^. Furthermore, specific analysis of the *Grn* KO mouse thalamus revealed a strong correlation between microglial GPNMB levels and lipofuscin (**Fig. 1j**), an undegradable autofluorescent material with variable lipid and protein composition that has been shown to accumulate in lysosomes during aging and neurodegeneration^38,39^, including in progranulin-deficient animals and NCL and FTD patients^40–42^. Overall, these data suggest that MG8, an emergent microglial population in the thalamus of aged *Grn* KO mice, is associated with lysosomal dysfunction in microglia.

### Epigenetic analysis of aged Grn KO microglia

To better understand the emergence of the diverse transcriptional states described above, we next assessed the enhancer landscape of microglia from aged mice. To do so, we purified microglial nuclei (PU.1^+^/NeuN^-^/OLIG2^-^) from 24-month WT and *Grn* KO mice (**Fig. 2a,b**) and performed bulk chromatin immunoprecipitation sequencing (ChIP-seq) after pulling down acetylated histone H3 lysine 27 chromatin (H3K27ac, a mark deposited by transcriptional coactivators with histone acetyltransferase activity^43^) to identify putative active enhancers in *Grn* KO microglia^43^. Analysis of H3K27ac signal at loci for cell type- specific markers confirmed successful enrichment for microglia (*P2ry12*, *Tmem119*, *Cx3cr1*) (**Fig. 2c**) and de-enrichment of other central nervous system cell types, including neurons (*Nefm*/*Nefl*), astrocytes (*Aldh1l1*), and oligodendroglia (*Olig1*/*Olig2*) (**Fig. 2d**). Thus, results of the bulk ChIP-seq reflect epigenetic changes specifically in microglia.

**Figure 2.**
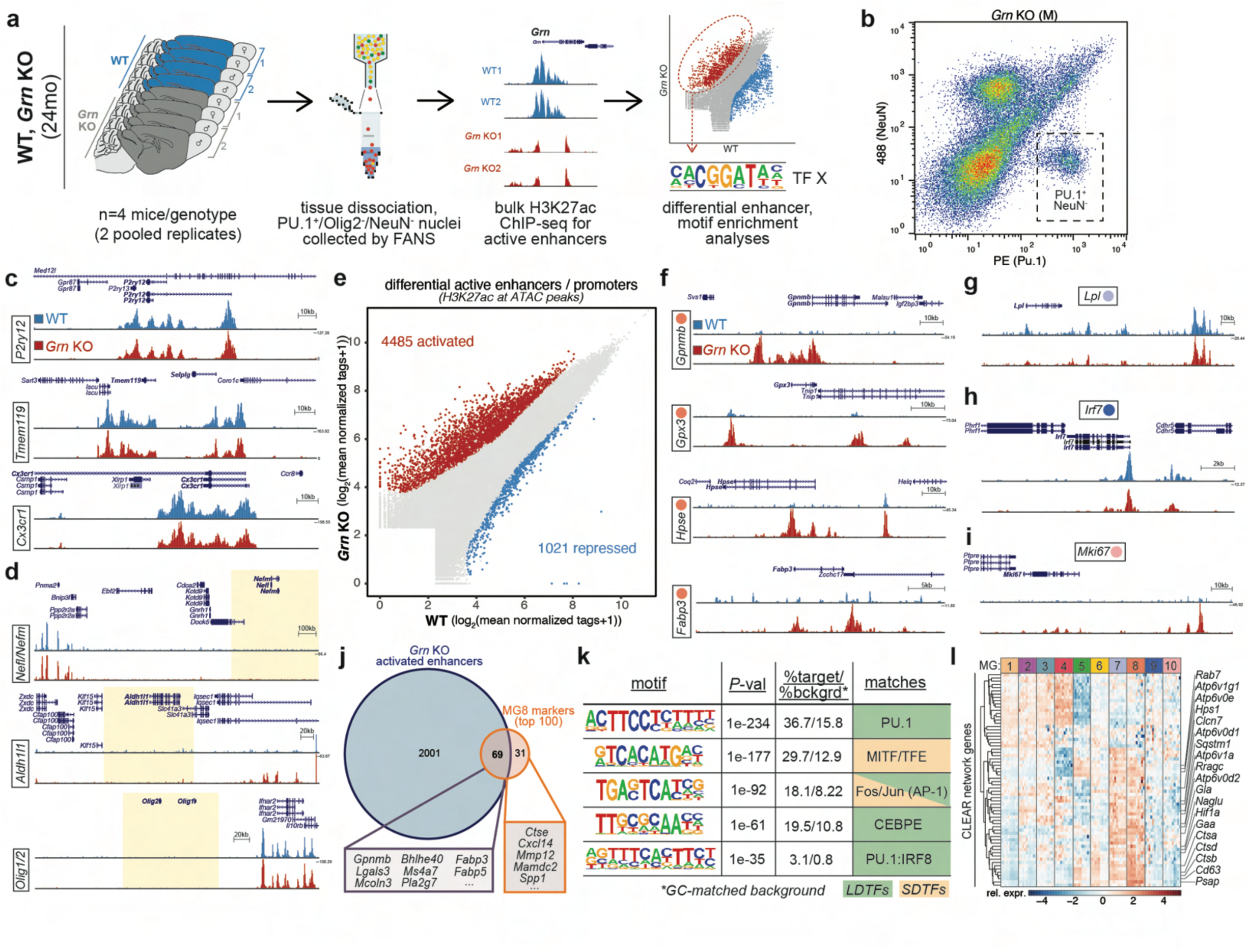
Analysis of active enhancers in aged *Grn* KO microglia. **(a)** Schematic outlining approach to enrich for PU.1^+^/NeuN^-^/Olig2^-^ microglia from aged (24- month) WT and *Grn* KO mouse brains for H3K27ac ChIP-seq (n=4 mice/genotype, 2 mice pooled per replicate). **(b)** Representative FACS plot displaying PU.1^+^/NeuN^-^ cell population collected for H3K27ac ChIP-seq. **(c)** Bulk H3K27ac signal at microglia-specific loci (*P2ry12*, *Tmem119*, *Cx3cr1*) in sorted cells to confirm successful enrichment for microglia. **(d)** Bulk H3K27ac signal at neuron-, astrocyte-, and oligodendroglia-specific loci (*Nefl/Nefm*, *Aldh1l1*, *Olig1/Olig2*, respectively) in sorted cells to confirm successful de-enrichment of non-microglial cells. Highlighted regions indicate proximal genomic regions of selected genes. **(e)** Scatter plot of differential H3K27ac activity at open chromatin between WT and *Grn* KO microglia. Red: regions with increased H3K27ac (FC>2), Blue: regions decreased H3K27ac tags (FDR<0.05). **(f)** Bulk H3K27ac signal at MG8-specific loci (*Gpnmb*, *Gpx3*, *Hpse*, *Fabp3*) in sorted cells, demonstrating emergent/higher peaks in *Grn* KO microglia. **(g)** Bulk H3K27ac signal at an MG7-specific locus (*Lpl*) in sorted cells, demonstrating lower peaks in *Grn* KO microglia. **(h)** Bulk H3K27ac signal at an MG9-specific locus (*Irf7*) in sorted cells, demonstrating lower peaks in *Grn* KO microglia. **(i)** Bulk H3K27ac signal at an MG10-specific locus (*Mki67*) in sorted cells, demonstrating emergent/higher peaks in *Grn* KO microglia. **(j)** Venn diagram displaying overlap of top 100 MG8 marker genes from scRNA-seq with activated enhancers, as determined by H3K27ac signal. **(k)** *De novo* motif enrichment analysis of activated enhancers in *Grn* KO microglia. **(l)** Relative expression of CLEAR network genes in microglia subclusters from scRNA-seq.

We then assessed differential active enhancer activity by comparing H3K27ac signal at regions of open chromatin across genotypes. This revealed significant increases and decreases of H3K27ac deposition (abs(FC>2), FDR<0.05) at 4,485 and 1,021 putative enhancer/promoter loci, respectively, in *Grn* KO microglia relative to age-matched WT microglia (**Fig. 2e and Supplementary Table 2**). Although a bulk microglial analysis, the differential H3K27ac signal captured changes in relative microglial subpopulation abundance across genotypes, as determined by scRNA-seq (**Fig. 1**), including increased signal at MG8-specific (*Gpnmb*, *Gpx3*, *Hpse*, *Fabp3*) and MG10- specific (*Mki67*) marker gene loci, and loss of signal at MG7-specific (*Lpl*) and MG9- specific (*Irf7*) marker loci (**Fig. 2f-2i**). Pathway analysis of nearest genes to active enhancers revealed enrichment in a variety of pathways related to core immune functions (GO biological process parent terms: positive regulation of locomotion, signaling by GPCR, intracellular signaling cassette, inflammatory response, regulation of cell activation, negative regulation of macrophage activation, etc.) and catabolic processes (GO biological process parent terms: negative regulation of lipid catabolic processes, positive regulation of hydrolase activity, regulation of lipid metabolic process, etc.), among others (**Extended Data** Fig. 2c **and Supplementary Table 2**).

Because MG8 was a microglial subpopulation exclusive to *Grn* KO mice (**Fig.1e- g**) that uniquely expressed a high number of marker genes not expressed in other microglial subpopulations (**Fig. 1d and Supplementary Table 1**), we next analyzed a large list of MG8-specific genes to determine if emergence of MG8 was associated with increased local putative enhancer activity. Among the top 100 unique marker genes expressed by MG8, 69 displayed significantly increased local H3K27ac signal in *Grn* KO microglia (**Fig. 2j**), providing evidence that the unique transcriptional states in *Grn* KO microglia result from histone modifications and alterations in transcription factor (TF) binding and activity. *De novo* motif analysis of all activated enhancer regions nominated several candidate lineage-determining TFs (LDTFs) and signal-dependent TFs (SDTFs) that likely collaborate to drive the diverse microglial states emergent in *Grn* KO mice (**Fig. 2k and Extended Data** Fig. 2d). Among these, motifs recognized by the LDTF PU.1 and the SDTF MITF/TFE TF family were the most enriched. The appearance of a PU.1 motif was expected and is consistent with its major role in driving the selection of microglia-specific enhancers through collaboration with additional microglia TFs. Of particular interest was the high enrichment of the MITF/TFE TF motif, suggesting the TF family’s likely role in driving aspects of gene expression in *Grn* KO microglia. Thus, we decided to focus subsequent analyses on this family of TFs.

Generally, members of the MITF/TFE family, comprised of basic helix-loop-helix-leucine zipper domain-containing TFs MITF, TFE3, TFEB, and TFEC, recognize a conserved, modified E-box motif found in the promoter regions of many autophagy and lysosome genes, collectively referred to as the coordinated lysosomal expression and regulation (CLEAR) network of genes^44,45^. Within our microglia scRNA-seq dataset, MG8 displayed the highest expression of the greatest number of CLEAR network genes (**Fig. 2l**). Additionally, expression of putative regulatory TFs across microglia subclusters revealed notable expression of *Mitf*, *Tfe3*, and *Tfeb* in MG8, with minimal expression of *Tfec* across all clusters (**Extended Data** Fig. 2e). However, because TF activity does not exclusively depend on high mRNA expression levels, but rather, appropriate nuclear localization, we next examined subcellular localization of MITF/TFE TFs in WT and *Grn* KO BMDMs, as well as BHLHE40 and BHLHE41, which have been shown to counteract MITF/TFE transcriptional activity^46,47^. Of these candidate TFs, only MITF and TFE3 displayed significant nuclear localization in *Grn* KO bone marrow-derived macrophages (BMDMs) (**Extended Data** Fig. 2f), suggesting these two TFs are likely to contribute to the transcriptional upregulation of key genes underlying the MG8 myeloid cell state. This notion is supported by findings from an accompanying manuscript (Balak *et al.*) in which ASO-mediated double knockdown of both *Mitf* and *Tfe3* mitigated the microglial transcriptional response in *Sgsh* KO mice, a model of mucopolysaccharidosis IIIA (MPS- IIIA), a different LSD. Collectively, these data demonstrate that MITF and TFE3 likely play a major role in coordinating the gene expression changes that underlie microglial states that emerge in response to lysosomal dysfunction.

### Cell autonomous alterations in Grn-deficient myeloid cells

Given the ability of microglia to sense their environment and respond to external stimuli, their high expression of *Grn* (especially in conditions of injury/CNS damage), and their high lysosomal function, whether the microglial states identified in *Grn* null mice arise strictly from a response to parenchymal damage/neurodegeneration or from cell autonomous deficits in microglia themselves remains to be determined. Conditional genetics approaches to deplete *Grn* in specific CNS cell populations in mice do not recapitulate the robust neuropathological deficits and transcriptional alterations observed in *Grn* null animals^48,49^, likely owing to adequate production and secretion by other neighboring cell types. Therefore, to interrogate the cell autonomous consequences of progranulin deficiency specifically in myeloid cells, we assessed transcriptional and functional changes in *Grn* KO mouse primary microglia (pMG) and BMDMs in isolation *in vitro*, expanding upon previous characterization of *Grn* KO myeloid cells^26,34,42,50–53^ (**Supplementary Table 3**).

Comparison of the top 150 upregulated genes in *Grn* KO pMG to our scRNA-seq dataset revealed their composite highest mean expression was most strongly associated with MG8 (**Fig. 3a**). Conversely, analysis of MG8-enriched marker genes revealed significant upregulation of nearly half of the top 175 MG8 markers, including many genes of interest related to lysosomes (*Atp6v0d2*, *Lgals3*, *Gpnmb*, etc.) and lipid processing (*Pla2g7*, *Pld3*, *Fabp5*, *Fabp3*, etc.), with less than 15% of MG8 marker genes being downregulated (**Fig. 3b,c**). Furthermore, *Grn* KO pMG displayed higher protein levels of key MG8-specific markers (GPNMB, FABP5, CatB), but not a MG7-specific marker (LPL) (**Fig. 3d,e**). Similar transcriptional and biochemical analyses in *Grn* KO BMDMs revealed consistent results (**Fig. 3f-j**). In addition to robust transcriptional alterations, isolated *Grn* KO BMDMs also displayed functional alterations, including increased proliferation (**Extended Data** Fig. 3a), altered phagocytosis (**Extended Data** Fig. 3b), and increased pro-inflammatory cytokine secretion (**Extended Data** Fig. 3c). The pronounced and conserved upregulation of a large set of MG8-specific markers in multiple *Grn* KO myeloid cell types suggests key features of an MG8-like state can be induced through cell autonomous deficits resulting from progranulin deficiency independent of neurodegeneration.

**Figure 3.**
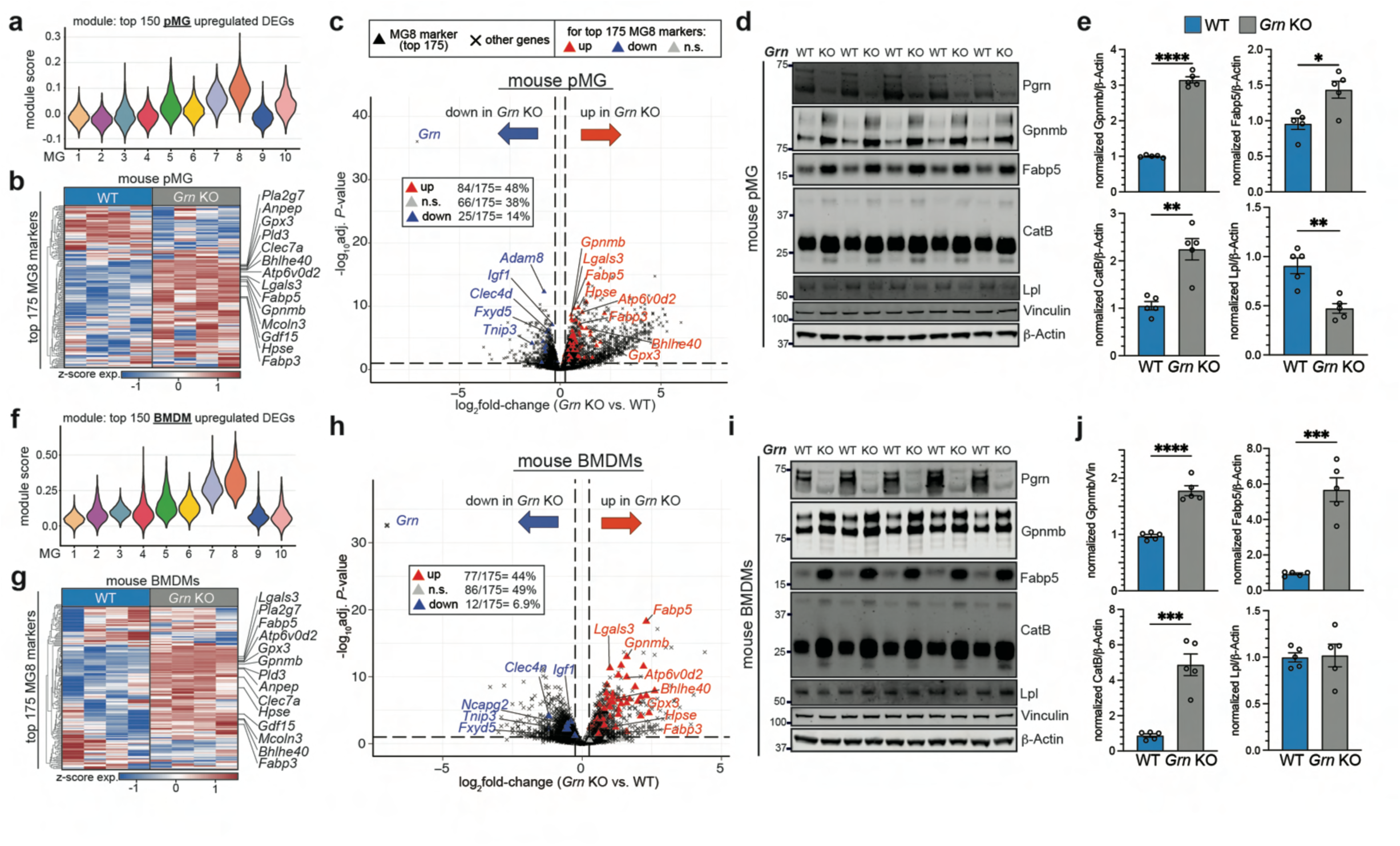
*Grn* KO myeloid cells display an MG8-like signature *in vitro*. **(a)** Average expression of top 150 regulated genes [from *Grn* KO primary microglia (pMG) *in vitro*] in each microglia subcluster from aged *Grn* KO microglia scRNA-seq. **(b)** Heatmap of relative expression of top 175 MG8 markers (from aged *Grn* KO microglia scRNA-seq) in WT and *Grn* KO pMG *in vitro*. **(c)** Volcano plot displaying differential expression between *Grn* KO and WT pMG. Top MG8 marker genes are highlighted as triangles. **(d,e)** Western blots and quantification of MG8 markers (GPNMB, FABP5, and total CatB) and LPL (not expressed by MG8) from protein lysates of WT and *Grn* KO mouse primary microglia (pMG) (n=5 biological replicates/genotype). **(f)** Average expression of top 150 upregulated genes [from *Grn* KO bone marrow-derived macrophages (BMDMs) *in vitro*] in each microglia subcluster from aged *Grn* KO microglia scRNA-seq. **(g)** Heatmap of relative expression of top 175 MG8 markers (from aged *Grn* KO microglia scRNA-seq) in WT and *Grn* KO BMDMs *in vitro*. **(h)** Volcano plot displaying differential expression between *Grn* KO and WT BMDMs. Top MG8 marker genes are highlighted as triangles. **(i)** Western blots and quantification of MG8 markers (GPNMB, FABP5, and total CatB) and LPL (not expressed by MG8) from protein lysates of WT and *Grn* KO mouse BMDMs (n=3 biological replicates/genotype). Data presented in (e,j) are mean±s.e.m. Student’s t-tests were performed to compare across genotypes in (e,j). \**P*<0.05, \*\**P*<0.01, \*\*\**P*<0.001, \*\*\*\**P*<0.0001.

Among the principal functions of macrophages and microglia, clearance of cellular debris via lysosome-mediated degradation is particularly important throughout aging and neurodegeneration. Lysosomes require the maintenance of several key properties (i.e. organellar pH, hydrolase activity, membrane integrity, macromolecular transport, etc.) to perform their physiological catabolic functions. Roles for progranulin and individual granulin peptides in regulating these diverse lysosomal properties have been identified, including in lysosomal lipid regulation^42,51,54–58^, cathepsin processing and function^59–62^, lysosomal pH^42,63^, and others. Because previous findings were obtained from a variety of tissues, primary cells, and cell lines, we aimed to comprehensively assess these lysosomal deficits using a single cellular system. Consistent with previous findings across various cell types, *Grn* KO BMDMs displayed decreased lysosomal proteolysis (**Extended Data** Fig. 3d), deacidified lysosomal pH (**Extended Data** Fig. 3e), reduced glucocerebrosidase activity (GCase) (**Extended Data** Fig. 3f), alterations in autophagy (**Extended Data** Fig. 3g), and lysosome membrane permeabilization (**Extended Data** Fig. 3h,i). These data highlight the utility of BMDMs for studying myeloid cell responses to lysosomal dysfunction.

### Identification of a conserved myeloid cell response to acute lysosomal stress

Because progranulin-deficient myeloid cells display deficits in numerous lysosomal properties, we next aimed to determine if any specific lysosomal impairments could explain the observed *Grn* KO myeloid cell phenotypes. To this end, we treated mouse BMDMs with a variety of pharmacological agents to differentially impact lysosomal function. We treated cells with Bafilomycin A1 (BafA1; v-ATPase inhibitor) to deacidify lysosomes, a cocktail containing E64, leupeptin, and pepstatin (E/L/P; protease inhibitors) to block hydrolytic activity of lysosomal cathepsins, U18666A (NPC1 inhibitor) to prevent lysosomal cholesterol egress, monosodium urate crystals (MSU; punctures phagosomal membranes) to induce lysosome membrane permeabilization (LMP), and VPS34IN1 (VPS34 inhibitor) to prevent endosome and autophagosome maturation/biogenesis by inhibiting phosphatidylinositol-3-phosphate synthesis (**Fig. 4a**). We first tested the response of BMDMs to increasing concentrations of each compound. Surprisingly, we observed an increase in cell number (nearly two-fold) after 22 hours of treatment with specific concentrations of each compound, except for MSU (**Fig. 4b**). We proceeded with the highest non-toxic concentration of each drug for subsequent experimentation (25nM BafA1, 6μM U18666A, 2μM VPS34IN1, 50μM E/L/P, 200μg/mL MSU). As expected, treatment with each compound at the selected dose disrupted the target lysosomal functions, including lysosomal pH (**Extended Data** Fig. 4a), lysosomal proteolysis (**Extended Data** Fig. 4b,c), lysosomal cholesterol egress (**Extended Data** Fig. 4d), endolysosomal vacuole size (**Extended Data** Fig. 4e,f), and lysosome membrane integrity (**Extended Data** Fig. 4g,h). These results confirmed that the selected non-toxic doses of each compound were also sufficient to impair lysosomal functions in BMDMs.

**Figure 4.**
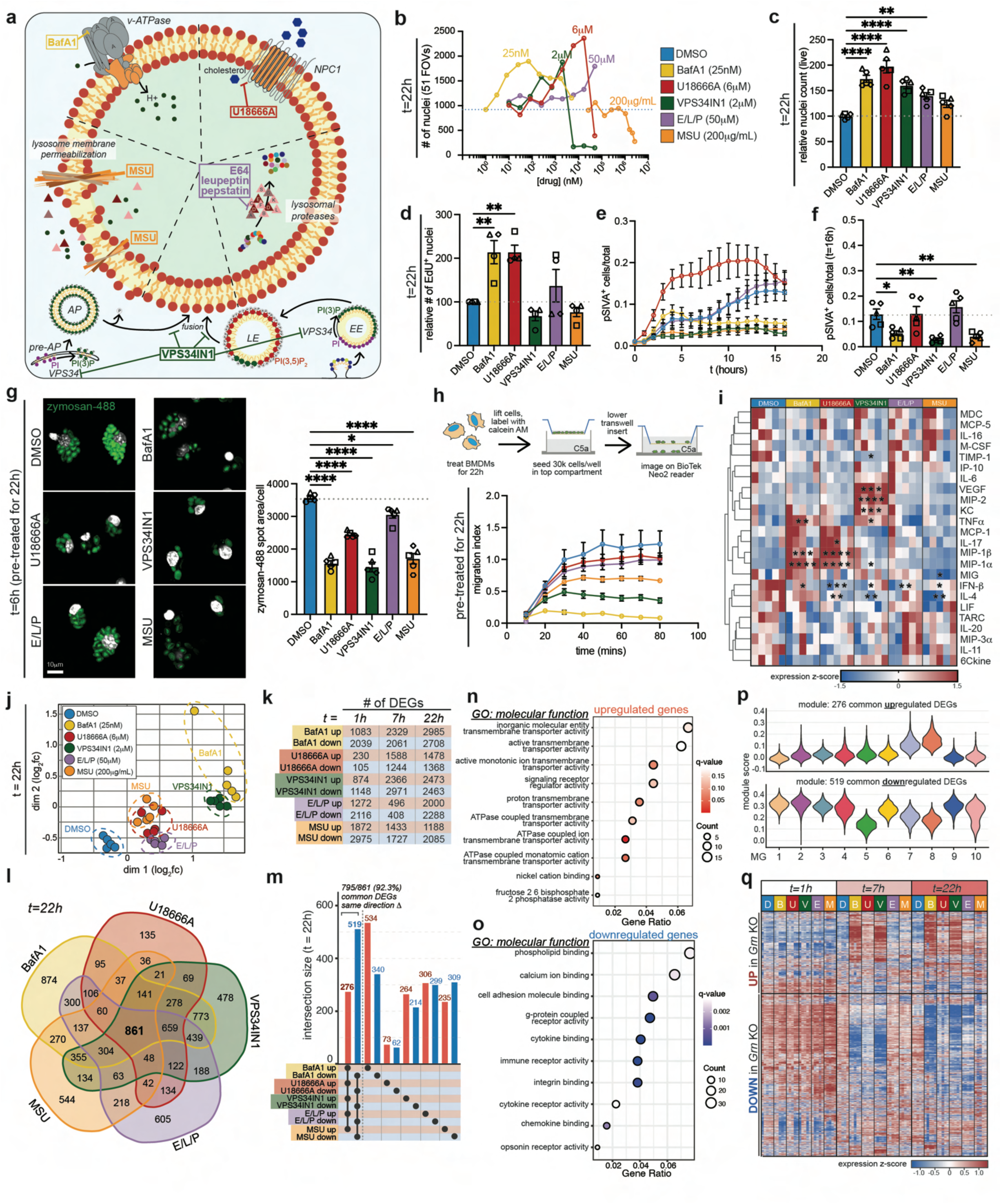
Pharmacological perturbation of lysosomal properties drives an acute functional and transcriptional remodeling of BMDMs. **(a)** Schematic displaying pharmacological approaches to perturb various lysosomal properties [BafA1, v-ATPase inhibitor; U18666A, NPC1 inhibitor; E64/leupeptin/pepstatin (E/L/P) cocktail, protease inhibitors; VPS34IN1, VPS34 inhibitor; monosodium urate (MSU), lysosome membrane permeabilizing agent]. **(b)** Dose response of WT BMDM live nuclei count in response to increasing doses of each lysosomal drug after 22 hours of treatment. Selected doses for downstream assays are displayed in the legend and above each curve (n=1 biological replicate/dose/drug). **(c)** Quantification of relative live nuclei count following a 22 hour treatment with various lysosomal drugs at selected doses (% of WT; n=5 biological replicates/genotype). **(d)** Quantification of EdU^+^ nuclei count following a 22 hour treatment with various lysosomal drugs at selected doses to measure proliferation (% of WT; n=4 biological replicates/genotype). **(e,f)** Quantification of pSIVA^+^ cells over time (e) and after 16 hours (f) of treatment with various lysosomal drugs at selected doses to measure transient phosphatidylserine exposure and apoptosis (fraction of total cells; n=5 biological replicates/genotype). **(g)** Representative images and quantification of zymosan-488 phagocytosis in WT BMDMs following a 22 hour pre-treatment with selected lysosomal drugs (n=5 biological replicates/genotype). **(h)** Schematic displaying experimental workflow and quantification to measure BMDM transwell migration towards chemoattractant C5a following a 22 hour pre-treatment with selected lysosomal drugs (n=5 biological replicates/genotype). **(i)** Heatmap displaying relative cytokine levels in conditioned media of BMDMs treated with lysosomal drugs for 22 hours (n=5 biological replicates/genotype). **(j)** Multidimensional scaling plot displaying overall transcriptional (dis)similarity between BMDMs treated with various lysosomal drugs for 22 hours. **(k)** Number of significant up- and downregulated differentially expressed genes (DEGs) over time in each condition (sampled at 1, 7, 22 hours). **(l)** Venn diagram displaying overlap in DEGs between conditions after 22 hours of treatment. **(m)** Bar graph displaying high degree of concordance in directionality of common DEGs between all treatment conditions (795/861 changed in same direction across all 5 treatments). **(n,o)** Gene ontology (GO: molecular function) analysis of (n) up- and (o) downregulated common genes. **(p)** Average expression of shared up- (top) and downregulated (bottom) shared DEGs in each microglia subcluster from aged *Grn* KO microglia scRNA-seq. **(q)** Heatmap displaying relative expression of constitutive *Grn* KO BMDM DEGs in BMDMs treated with lysosomal drugs for 22 hours (n=5 biological replicates/genotype). Data presented in (c-h) are mean±s.e.m. One-way ANOVA tests with multiple comparisons were performed to compare across treatment conditions in (c,d,f,g). \**P*<0.05, \*\**P*<0.01, \*\*\**P*<0.001, \*\*\*\**P*<0.0001.

Next, we profiled the acute functional alterations in response to the pharmacological lysosomal perturbations. Consistent with our previous dosing experiment (**Fig. 4b**), treatment with all compounds, except MSU, resulted in an increase in live cell number (**Fig. 4c**), which was attributed to increased proliferation (BafA1, U18666A) (**Fig. 4d**) and/or increased survival (BafA1, VPS34IN1, MSU) (**Fig. 4e,f**). Treatment with any of the lysosomal drugs resulted in reduced phagocytosis (**Fig. 4g**) and C5a-mediated chemotaxis (**Fig. 4h**). Additionally, pharmacological perturbations of the lysosome impacted cytokine secretion, generally favoring an increase in secretion of pro-inflammatory chemokines/cytokines including MIP-1α, MIP-1β, and TNFα, among others, while reducing IFN-β secretion (**Fig. 4i**). Collectively, these data demonstrate that directly disrupting lysosomal homeostasis is sufficient to impact major myeloid cell functions.

To obtain a more comprehensive view of how diverse lysosomal perturbations impact myeloid cell states, we performed bulk RNA-seq to assess transcriptome-wide changes after 1, 7, and 22 hours of treatment. Time series data demonstrated a progressive transcriptional response of BMDMs to drug treatment (**Supplementary Table 4**), with all treatment groups displaying large transcriptional changes relative to DMSO vehicle-treated cells by 22 hours of treatment (**Fig. 4j,k**). While all groups displayed many significantly differentially expressed genes (DEGs) by 22 hours, a large number of overlapping DEGs across all treatments was observed (**Fig. 4l**). Of the 861 genes that were significantly differentially expressed in all conditions, 795 of the 861 (92.3%) genes were changed in the same direction across all conditions (**Fig. 4m, Supplementary Table 4**). In other words, this set of genes was commonly up- or downregulated across all five treatment groups (with minimal diverging directionality) and therefore, represents a core myeloid cell transcriptional response to lysosomal stress.

Unsurprisingly, pathway analysis of the common upregulated genes highlighted pathways related to ATPase coupled ion transmembrane transporter activity, consistent with a homeostatic compensatory induction of v-ATPase-related genes in response to lysosomal stress (**Fig. 4n**). On the other hand, pathway analysis of downregulated genes revealed broad changes related to cytokine/chemokine binding and immune receptor activity, as well as cell adhesion molecule and integrin binding (**Fig. 4o**), which may explain the functional alterations in phagocytosis and chemotaxis upon lysosomal perturbation. Intriguingly, besides many immune-related genes, several genes that modify AD risk, including *Pilra*, *Inpp5d*, *Sorl1*, *Abca1*, *Picalm*, *Adam10*, *Sppl2a*, *Maf*, and *Abi3*, were among the downregulated DEGs shared across all lysosomal perturbations (**Supplementary Table 4**). In addition to the shared DEGs highlighted above, each specific perturbation also led to sets of distinct gene expression changes (**Supplementary Table 4**), demonstrating that unique nuances in myeloid cell transcriptional states can emerge in response to lysosomal damage depending on the precise nature of the dysfunction. However, ruling out extralysosomal effects of these perturbations is necessary to determine which specific changes are attributed solely to disruption to a particular lysosomal property.

To gain insights into putative TFs driving treatment-dependent changes in gene expression, we performed *de novo* motif enrichment analysis of the promoter regions residing -2kb to +500bp from the transcriptional start sites of the upregulated genes for each condition (FDR<0.05) and for the shared set of 276 genes upregulated in all conditions. This promoter biased analysis will not capture distal signal-dependent enhancers or LDTFs but has the potential to identify motifs for signal-dependent factors that act at both enhancers and promoters. Notably, a consensus binding site for MITF/TFE factors was the sole significantly enriched motif in the promoter regions of the 276 shared upregulated genes (**Extended Data** Fig. 5a). In accordance with this finding, variations of consensus MITF/TFE motifs were among the top three most highly enriched motifs in responsive promoter regions of each of the individual treatment conditions, with the remaining enriched motifs corresponding to binding sites for general promoter transcription factors (**Extended Data** Fig. 5b-e). These findings are consistent with major roles of MITF/TFE factors in driving the altered programs of gene expression not only in *Grn* KO microglia as inferred from the enhancer analysis described above, but also in pharmacologically-induced lysosomal dysfunction in BMDMs.

Because *TREM2* is required for the microglial transition out of a homeostatic state in certain contexts^10,11,51,64^, we next assessed whether *Trem2* was involved in the myeloid cell transcriptional response to lysosomal dysfunction (**Extended Data** Fig. 6a**, Supplementary Table 5**). Of the DEGs that were significantly changed upon BafA1 treatment in WT but not in *Trem2* KO BMDMs, many genes were already altered in the same direction in DMSO-treated *Trem2* KO cells relative to WT (**Extended Data** Fig. 6b,c), suggesting basal lysosomal impairments could result from *Trem2* deficiency, consistent with a recent study in *TREM2* KO human microglia^65^. Nonetheless, over 3,000 BafA1-induced DEGs were similarly changed in *Trem2* KO BMDMs treated with BafA1 (**Extended Data** Fig. 6b,d), including nearly all of the 795 DEGs commonly changed across the diverse pharmacological perturbations (**Extended Data** Fig. 6e). *Trem2* KO BMDMs treated with BafA1 also increased production of pro-inflammatory chemokines/cytokines MIP-1α, MIP-1β, and TNFα (**Extended Data** Fig. 6f). Furthermore, the transcriptional and cytokine responses of BMDMs to lysosomal dysfunction were largely conserved in pMG treated with BafA1 (**Extended Data** Fig. 7a**- d and Supplementary Table 6**), suggesting these common changes likely represent a core set of genes that defines a myeloid cell-specific acute lysosomal stress response. Interestingly, among the microglial subpopulations identified in our scRNA-seq dataset, the shared upregulated genes were the most enriched in MG8, the aged *Grn* KO-specific microglial population, while the downregulated genes were de-enriched in this population, providing additional evidence that MG8 likely emerges due to lysosomal dysfunction (**Fig. 4p**). Finally, analysis of DEGs from constitutive *Grn* KO BMDMs (**Fig. 3**) revealed the greatest overlap with the BMDM response to BafA1 and VPS34IN1 treatment (**Fig. 4q**), suggesting *Grn* KO phenotypes may be driven by lysosomal pH deficits or changes in endosome and/or autophagosome maturation. Collectively, these data demonstrate that primary lysosomal dysfunction is sufficient to drive a TREM2-independent transcriptional and functional remodeling of myeloid cells characterized by increased expression of lysosome-related genes, secretion of pro-inflammatory cytokines, reduction of phagocytosis and migration, and downregulation of many immune-related genes, including several AD risk genes enriched in microglia.

### Lysosomal deacidification contributes to Grn KO phenotypes

As pharmacological perturbations induce rapid, acute, and often sustained effects due to irreversible binding/inhibition of its targets, we next sought to determine the effect of a more progressive disruption of lysosomal function by using a genetic targeting approach. To do this, we transfected WT BMDMs with ribonucleoproteins complexes (RNPs) containing SpCas9 and single-guide RNAs (sgRNAs) targeting the genes encoding proteins that regulate key lysosomal properties (**Fig. 5a**)^66^. Genes were selected to target analogous lysosomal properties to those modulated in the pharmacological experiments above, including *Rosa26* (vehicle, positive cutting control), *Atp6v1a* (BafA1), *Npc1* (U18666A), *Pik3c3* (VPS34IN1), *Ctsd* (E/L/P), and these results were compared directly with cells in which *Grn* or *Trem2* were knocked out using a similar approach. This arrayed approach resulted in the rapid mutagenesis of the target loci and the prevention of *de novo* transcription of full-length target mRNAs. We confirmed highly efficient reduction of target proteins over the course of two weeks, with greater than 85% of protein lost by day 6 following transfection for all targets (**Fig. 5b,c**). Therefore, in addition to providing a model to study the BMDM response to progressive loss of the protein of interest, this approach also enabled us to examine the longer-term homeostatic adaptations to persistent lysosomal stress following sustained loss of each target protein.

**Figure 5.**
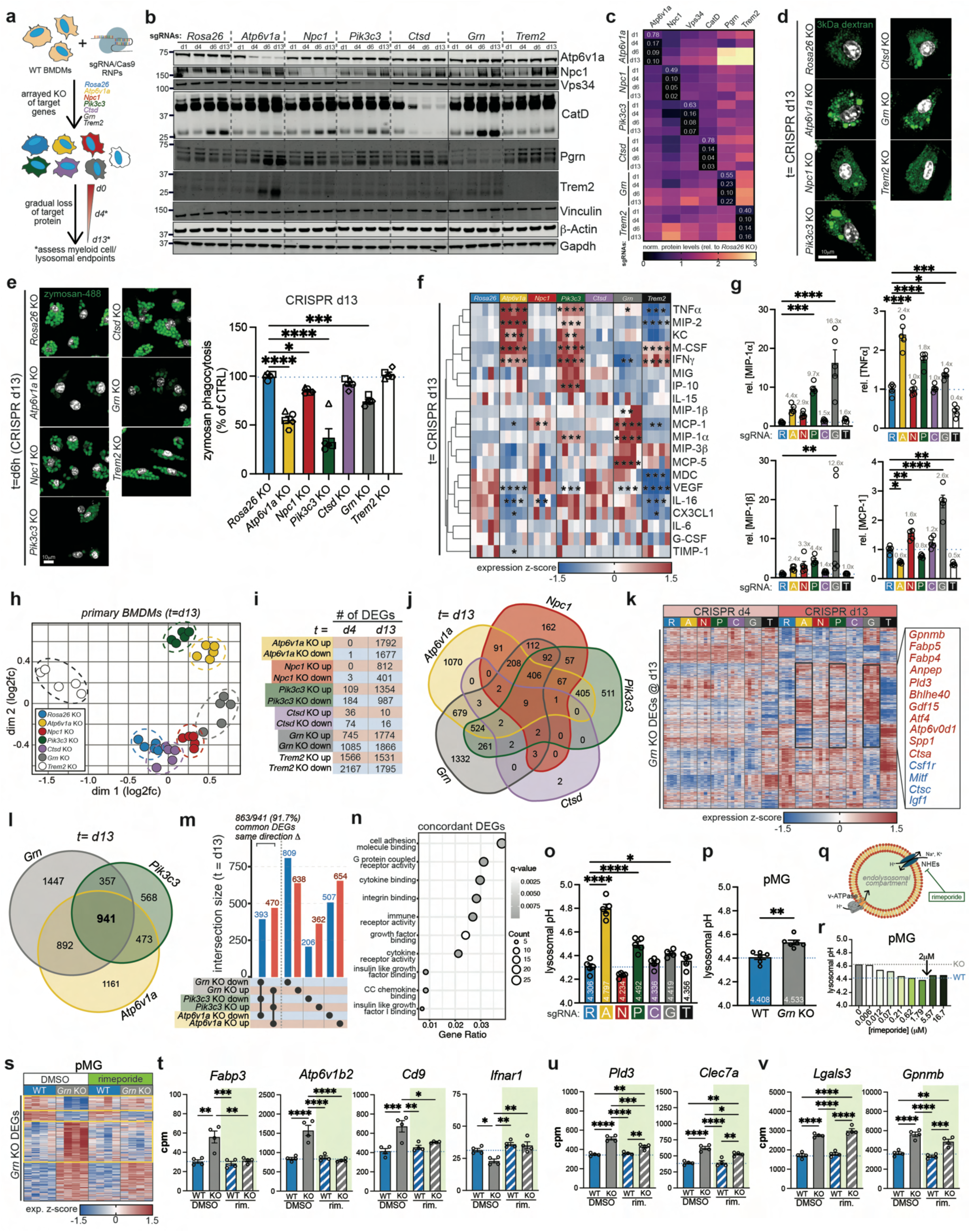
Genetic perturbation of lysosomal properties drives a progressive functional and transcriptional remodeling of BMDMs. **(a)** Schematic displaying approach to genetically perturb various lysosomal properties using CRISPR/Cas9 ribonucleoprotein (RNP) delivery in WT BMDMs. **(b,c)** Western blots (b) and quantification (c) displaying progressive loss of target proteins following transfection with sgRNA/Cas9 RNPs relative to *Rosa26* KO cells. **(d)** Representative images of BMDMs 22 hours after loaded with fluorescent 3 kDa dextran at d13 post-CRISPR. **(e)** Representative images and quantification of zymosan-488 phagocytosis in BMDMs following at d13 post-CRISPR (n=5 biological replicates/genotype). **(f,g)** Heatmap displaying relative cytokine levels in conditioned media of BMDMs at d13 post- CRISPR (n=5 biological replicates/genotype). **(h)** Multidimensional scaling plot displaying overall transcriptional (dis)similarity between BMDMs at d13 post-CRISPR. **(i)** Number of significant up- and downregulated differentially expressed genes (DEGs) over time in each condition (sampled at d4 and d13 post-CRISPR). **(j)** Venn diagram displaying overlap in DEGs between conditions at d13 post-CRISPR. **(k)** Heatmap displaying relative expression of constitutive *Grn* KO BMDM DEGs in BMDMs at d4 and d13 post-CRISPR (n=4-5 biological replicates/genotype). **(l)** Venn diagram displaying overlap in DEGs between *Atp6v1a*, *Pik3c3*, and *Grn* KO BMDMs at d13 post-CRISPR. **(m)** Bar graph displaying high concordance in directionality of common DEGs between *Atp6v1a*, *Pik3c3*, and *Grn* KO BMDMs (863/941 changed in same direction across all 3 genotypes). **(n)** Gene ontology (GO: molecular function) analysis of all concordant DEGs shared by *Atp6v1a*, *Pik3c3*, and *Grn* KO BMDMs. **(o,p)** Measurement of lysosomal pH in (o) BMDMs at d13 post-CRISPR and (p) WT and *Grn* KO pMG (n=4-5 biological replicates/genotype). **(q)** Schematic displaying strategy to acidify endolysosomes using sodium/proton exchanger (NHE) inhibitor rimeporide. **(r)** Measurement of lysosomal pH in *Grn* KO pMG following 72 hour treatment with rimeporide (n=1 biological replicates/dose). **(s)** Heatmap displaying relative expression of constitutive *Grn* KO pMG DEGs with and without treatment with 2μM rimeporide for 72 hours (n=4 biological replicates/genotype). **(t-v)** Normalized expression of *Grn* KO DEGs of interest that were corrected (t), partially corrected (u), or unaffected (v) by 72 hour rimeporide treatment of pMGs (n=4 biological replicates/genotype). Data presented in (e,g,o,p,t-v) are mean±s.e.m. One-way ANOVA tests with multiple comparisons were performed to compare across treatment conditions in (e,g,o,t-v). Student’s t- tests were performed to compare across genotypes in (p) \**P*<0.05, \*\**P*<0.01, \*\*\**P*<0.001, \*\*\*\**P*<0.0001.

Because over 90% of target protein was missing for an extended duration (**Fig. 5c**) and alterations in endolysosomal morphology were observed by day 13 (**Fig. 5d**), we assessed several myeloid cell phenotypes of interest 13 days post-CRISPR. Similar to the acute pharmacological lysosomal treatments, genetic perturbations to key lysosomal genes resulted in reduced zymosan phagocytosis in all conditions except the *Ctsd* KO BMDMs (**Fig. 5e**). In this context, *Trem2* KO BMDMs also displayed comparable phagocytosis to control cells (**Fig. 5e**), potentially due to high (50ng/mL) M-CSF supplementation, which we have previously shown can rescue phagocytosis impairments associated with TREM2 deficiency^64^. Although supernatant from *Atp6v1a* and *Pik3c3* KO BMDMs contained significantly higher M-CSF (**Fig. 5f**), this was not sufficient to rescue phagocytosis deficits, suggesting this impairment downstream of lysosomal dysfunction is either CSF1R- and TREM2-independent or that lysosomal dysfunction reduces sensitivity to M-CSF stimulation. Furthermore, secretion of several proinflammatory cytokines/chemokines (TNFα, MIP-1α, MIP-1β) by *Atp6v1a* and *Pik3c3* KO BMDMs most closely resembled *Grn* KO BMDMs (**Fig. 5f,g**).

Next, to comprehensively evaluate the acute and compensatory transcriptional responses of BMDMs to genetic lysosomal perturbations, we performed bulk RNA-seq at day 4 and day 13 post-CRISPR KO of the same genes described above (**Fig. 5h**). Although target protein loss by day 4 was greater than 75% in all conditions (**Fig. 5b,c**), only some genotypes displayed substantial differential gene expression at this timepoint (*Grn* KO = 1,830 DEGs; *Trem2* KO = 3,733 DEGs), while others did not (*Atp6va1* KO = 1 DEG; *Npc1* KO = 3 DEGs; *Pik3c3* KO = 293 DEGs; *Ctsd* KO = 110 DEGs) (**Fig. 5i and Supplementary Table 7**). However, by day 13, all genotypes displayed a large number of DEGs (*Atp6va1* KO = 3,469 DEG; *Npc1* KO = 1,213 DEGs; *Pik3c3* KO = 2,341 DEGs; *Grn* KO = 3,640 DEGs; *Trem2* KO = 3,326 DEGs), except *Ctsd* KO, which displayed even fewer DEGs than at day 4 (26 DEGs at day 13), suggesting that BMDMs can engage adequate compensatory mechanisms to mitigate the loss of Cathepsin D and prevent lysosomal dysfunction during the assessed experimental timeframe. Across all genotypes, *Atp6v1a* and *Pik3c3* KO shared the greatest overlap with *Grn* KO BMDMs, which included many genes that were also upregulated in *Grn* KO MG8 compared to homeostatic microglia from our scRNA-seq dataset, as well as in *Grn* KO pMG, including *Gpnmb*, *Fabp5*, *Anpep*, *Pld3*, *Bhlhe40*, *Gdf15*, and others (**Fig. 5j,k and Supplementary Tables 1,3,7**). Among the 941 DEGs shared between *Atp6v1a*, *Pik3c3*, and *Grn* KO BMDMs (**Fig. 5l**), 863 genes (91.7%) were altered in concordant directions across all three genotypes (**Fig. 5m**). This concordant gene set was enriched in DEGs related to cell adhesion (GO terms: cell adhesion molecular binding, integrin binding, etc.) and various immune functions (GO terms: cytokine binding, immune receptor activity, cytokine receptor activity, etc.) (**Fig. 5n**). Overall, these results highlight the shared transcriptional and functional alterations resulting from sustained loss of *Grn*, *Atp6v1a*, and *Pik3c3* and support the notion that *Grn* KO myeloid cell phenotypes may involve similar cellular pathways to those affected by inhibition/loss of v-ATPase/*Atp6v1a* and VPS34/*Pik3c3*.

As progranulin has pleiotropic lysosomal functions (**Extended Data** Fig. 3), we used the results of our targeted pharmacological and genetic perturbations to narrow down key pathways that could underlie the observed *Grn* KO myeloid cell changes. Interestingly, genetic KO of *Atp6v1a*, *Pik3c3*, or *Grn* in BMDMs all resulted in lysosomal deacidification (**Fig. 5o**), which was also observed in BafA1- or VPS34IN1-treated BMDMs (**Extended Data** Fig. 4a), constitutive *Grn* KO BMDMs (**Extended Data** Fig. 3e), and constitutive *Grn* KO pMG (**Fig. 5p**). Pharmacological sodium/proton exchanger (NHE) inhibition with rimeporide^67^ (**Fig. 5q**) promoted a dose-dependent endolysosomal reacidification in *Grn* KO pMG, which was restored to WT levels at 2μM (**Fig. 5r**). Importantly, 72 hour rimeporide treatment corrected more than half of the transcriptional alterations of *Grn* KO pMG (**Fig. 5s and Supplementary Table 8**), with many DEGs being fully (**Fig. 5t**) or partially corrected (**Fig. 5u**), while others were unaffected (**Fig. 5v**). Additionally, because v-ATPase or VPS34 inhibition can also affect autophagy^68,69^, which has been shown to be defective in *Grn* KO cells^70^ (**Extended Data** Fig. 3g), we assessed the consequences of genetically ablating key components of the canonical macroautophagy machinery, including *Atg7* (LC3 lipidation), *Atg14* (autophagosome formation), and *Uvrag* (autophagosome formation) (**Extended Data** Fig. 8a,b). Although target knockdown was highly efficient (**Extended Data** Fig. 8c) and predicted effects on autophagosome maturation were observed (**Extended Data** Fig. 8d,e), these conditions did not phenocopy the GPNMB increase or cytokine secretion changes observed in *Atp6v1a*, *Pik3c3*, and *Grn* KO BMDMs (**Extended Data** Fig. 8d-f). Together, these data demonstrate that lysosomal deacidification, and not canonical macroautophagy impairments, likely underlie a large proportion of phenotypes seen in *Grn* KO myeloid cells.

### Gpnmb upregulation upon lysosomal stress promotes acidification

In recent years, GPNMB has emerged as a potential biomarker and/or therapeutic target for several central and peripheral indications, including PD^71,72^, obesity^73^, and various cancers^74–76^, among others. Because *Gpnmb* was among the top unique and defining markers of MG8, the *Grn* KO-specific microglial population (**Fig. 1h and 6a**), and positively correlated with autofluorescent lipofuscin accumulation in aged microglia (**Fig. 1l and 6b**), we sought to better understand its role in myeloid cells, especially in the context of progranulin deficiency and lysosomal homeostasis. Across all different *Grn* KO systems tested, including constitutive and CRISPR KO BMDMs, pMG, and microglia from aged mice, *Gpnmb* mRNA and protein levels were significantly elevated (**Fig. 1k, 3, 5k, 6c, Extended Data** Fig. 8d**, and Supplementary Tables 1, 3, 7**). Interestingly, pharmacological treatment of BMDMs with BafA1 led to a massive redistribution of a perinuclear GPNMB pool into lysosomes, as detected by super resolution microscopy (**Fig. 6d**). Additionally, treatment with all selected lysosomal drugs resulted in an increase of *Gpnmb* mRNA and protein relative to DMSO vehicle-treated cells, which was not observed in BMDMs treated with lipopolysaccharide (LPS) or AZD8055 (mTOR inhibitor) (**Fig. 6e,f**). Furthermore, genetic perturbation of *Atp6v1a*, *Npc1*, *Pik3c3*, or *Grn* resulted in a time-dependent increase in expression of *Gpnmb* (**Fig. 6g**). These data demonstrate that *Gpnmb* is transcriptionally and post-translationally elevated in myeloid cells in response to diverse lysosomal stress that is distinct from activation that occurs through LPS or mTOR inhibition.

**Figure 6.**
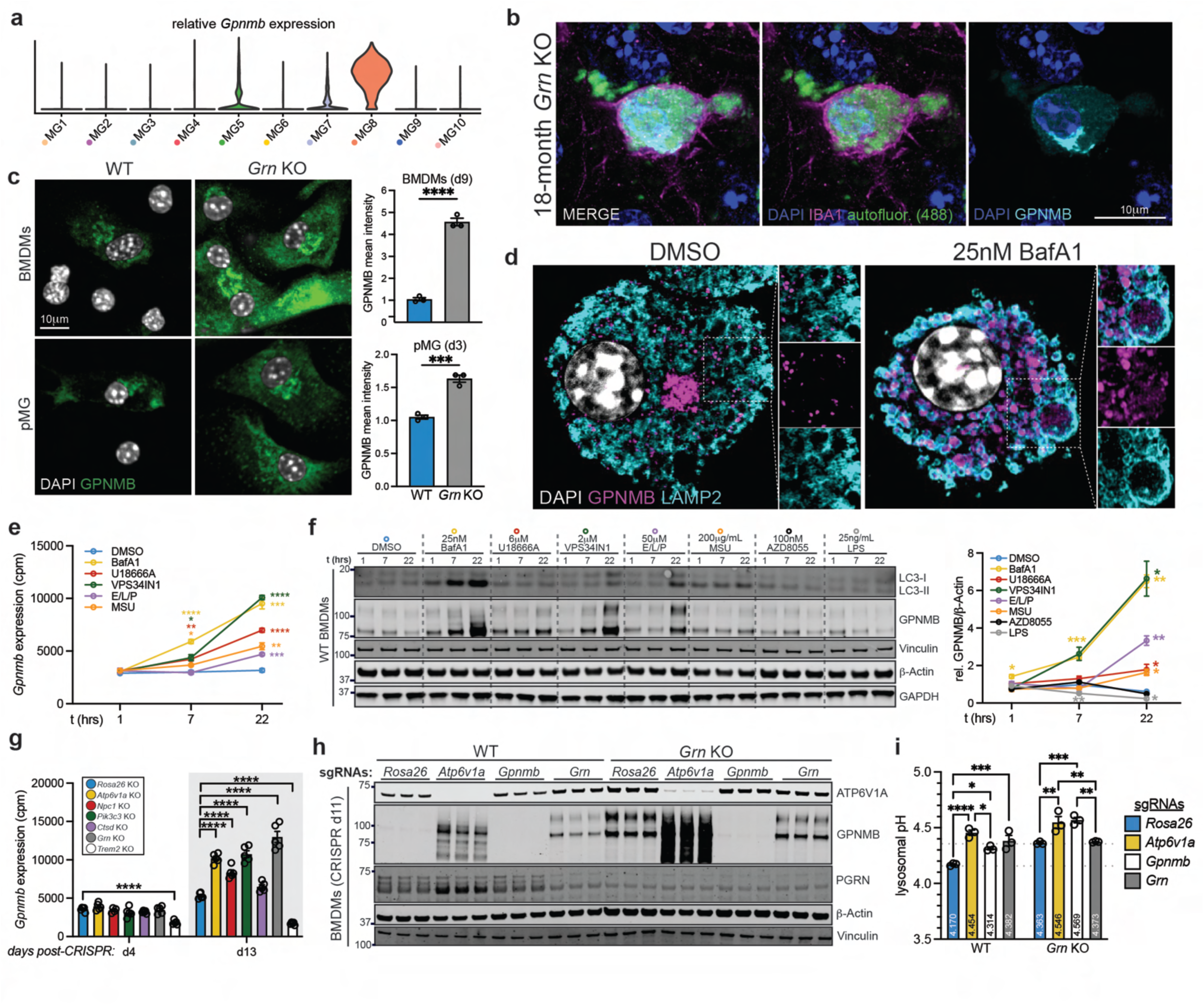
GPNMB is upregulated upon lysosomal perturbations and promotes reacidification. **(a)** Violin plot displaying MG8-restricted expression of *Gpnmb* in aged *Grn* KO scRNA-seq microglia subclusters. **(b)** Super-resolution representative images of lipofuscin-laden GPNMB-positive microglia in aged *Grn* KO mouse. **(c)** Representative images and quantification of GPNMB immunostaining in WT and *Grn* KO BMDMs (top) and pMG (bottom) (n=3 biological replicates/genotype). **(d)** Representative super-resolution image of WT BMDMs treated with DMSO or BafA1 for 22 hours, displaying lysosomal localization of GPNMB upon BafA1 treatment. **(e,f)** *Gpnmb* mRNA (e) and protein (f) expression over time following treatment with various lysosomal drugs (BafA1, U18666A, VPS34IN1, E/L/P, MSU), mTOR inhibitor (AZD8055), and LPS (e, n=5 biological replicates/condition; f, n=4 biological replicates/condition). **(g)** *Gpnmb* mRNA expression over time following CRISPR/Cas9-mediated knockout of various genes of interest (n=5 biological replicates/genotype). **(h)** Western blot demonstrating successful knockout of ATP6V1A, GPNMB, and PGRN using CRISPR/Cas9 in WT and *Grn* KO BMDMs. **(i)** Measurement of lysosomal pH in compound loss-of-function BMDMs at d11 post-CRISPR (n=3 biological replicates/condition). Data presented in (c,e-g,i) are mean±s.e.m. One-way ANOVA tests with multiple comparisons were performed to compare across treatment conditions at each timepoint in (e-g,i). \**P*<0.05, \*\**P*<0.01, \*\*\**P*<0.001, \*\*\*\**P*<0.0001.

GPNMB acts as an anti-inflammatory signaling molecule via numerous extracellular mechanisms, including its cleavage, secretion, and engagement with cell surface receptors on surrounding cells^77–79^. On the other hand, its intracellular functions are less characterized. Because we observed an upregulation and lysosomal translocation of GPNMB upon lysosomal stress (**Fig. 6d**), and GPNMB has been previously reported to chaperone v-ATPase assembly in senescent endothelial cells^80^, we examined the effect of modulating GPNMB on lysosomal pH in WT and *Grn* KO cells. In WT BMDMs, KO of *Gpnmb* resulted in lysosomal deacidification to a level comparable to *Grn* KO cells (**Fig. 6h,i**). Consistent with previous results (**Fig. 5o and Extended Data** Fig. 3e), *Grn* KO BMDMs displayed increased GPNMB protein and lysosomal deacidification, and KO of *Gpnmb* further increased lysosomal pH to the same extent as *Atp6v1a* KO BMDMs on the same genetic background (**Fig. 6h,i**). Thus, GPNMB is required for the homeostatic response to promote lysosomal acidification upon lysosomal damage, which can indirectly contribute to its anti-inflammatory effects.

## DISCUSSION

In this study, we have identified an intimate link between lysosomal health and myeloid cell states, in which myeloid cells respond to acute and/or chronic lysosomal insults and undergo robust transcriptional, epigenetic, and functional remodeling. We describe the transcriptional response (comprised of nearly 600 DEGs with consistent directionality) that murine macrophages and primary microglia undergo upon pharmacologic disruption of diverse lysosomal properties. This shared signature, which we believe constitutes a generalized myeloid cell response to lysosomal damage, is characterized by altered expression of key lysosome and immune-related genes and is associated with the numerous functional alterations we observed. Importantly, much of this transcriptional signature is represented in adult microglia from multiple animal models of neurodegenerative and LSDs, including progranulin-deficient mice and *Sgsh* mutant mice (accompanying paper, Balak *et al.*). Along similar lines, our analyses (this manuscript and Balak *et al.*) of enhancer activity from microglia isolated from mouse brain nominated a common set of transcription factors as mediators of the disease-associated emergent microglial subpopulations from these two disease models, further supporting the notion of a core response of myeloid cells to lysosomal dysfunction.

Overall, our data are consistent with a model whereby overwhelmed lysosomes signal to promote a variety of transcriptional, and ultimately functional, responses leading to the expansion of the myeloid cell pool via proliferation/chemokine secretion and possibly survival to promote resolution of inflammation and clearance of parenchyma damage. This contrasts with responses typically observed in non-myeloid cells, where decreased lysosomal health generally reduces cell viability. If lysosomal dysfunction is chronic, as in the case of progranulin deficiency, reactive microglia may become deleterious. Due to their secretion of pro-inflammatory cytokines/chemokines, along with their reduced phagocytic and lysosomal degradative capacity, we speculate that they likely exacerbate pathology in the context of neurodegeneration, possibly comprising the neurotoxic microglia that can induce TDP-43 pathology in progranulin-deficient mice^26^ or after concomitant ablation of *Grn* and *Tmem106b*, a genetic modifier of FTD-*GRN*^35^; however, it remains to be conclusively determined if these lysosomal stress-associated microglia are functionally beneficial or are a maladaptive and detrimental population in the brain. By leveraging the unique markers we identified here, follow-up studies can begin to experimentally dissect the role of these specific microglial subpopulations in disease pathology and progression, which may inform drug development efforts aimed at curbing neuroinflammation across multiple neurodegenerative diseases and LSDs.

Secreted progranulin has been previously described to exert anti-inflammatory effects primarily through interactions with TNFRs^81^, TLR9^28^, sPLA2-IIA^82^, and other signaling pathways^83,84^. Our data support an additional indirect mechanism through which progranulin suppresses inflammation—namely, via its role in maintaining proper lysosomal health, as targeted perturbations to various lysosomal properties regulated by progranulin were sufficient to drive immune responses in macrophages and microglia. We aimed to use the results of our targeted perturbation studies to better understand how impairments in specific lysosomal properties can contribute to the phenotypes seen in progranulin-deficient myeloid cells. To this end, we observed the highest overlap in transcriptional and functional outcomes between *Grn* KO cells and conditions that induced lysosomal deacidification, such as pharmacologic or genetic disruption of the lysosomal v-ATPase or VPS34. Furthermore, endolysosomal reacidification using rimeporide, an inhibitor of sodium/proton exchangers including those present in the endolysosomal system, rescued more than half of the gene expression changes observed in *Grn* KO pMG, highlighting lysosomal pH as a major driver of *Grn* KO-associated phenotypes despite the fact the lysosomal deacidification is rather subtle in this disease model. It is important to note that individual lysosomal properties function interdependently and can cross-regulate each other. Therefore, because many hydrolases, channels, and other lysosomal proteins are highly pH-sensitive, small changes in lysosomal pH are likely to affect many aspects of lysosomal biology. Whether the pH-sensitive changes in *Grn* KO cells are due to signaling processes directly downstream of lysosomal deacidification, or due to secondary effects on other pH-dependent lysosomal processes, remains an open question. Nonetheless, these findings emphasize the necessity of a tight regulation of lysosomal pH for suppressing myeloid cell reactivity and inflammation.

Lysosomes have a remarkable ability to sense changes in macromolecular content and other organellar properties, activating a variety of mechanisms to restore normal function when altered. Among these, the transcriptional induction of CLEAR motif-containing genes via MITF/TFE TFs is a well-conserved process active in most cell types to promote autophagosome and lysosome biogenesis^44,45^. In myeloid cells, activation of these TFs increases lysosomal gene expression and also coordinates immune responses, as shown here and previously in response to LPS^85^, certain bacterial exposures^86,87^, and changes in lysosomal hydrostatic pressure^88^. The precise gene expression programs that are engaged likely depend on differential enhancer activation and on the cooperative binding with divergent TFs that are co-expressed in these diverse contexts^89^. With respect to the data we presented here, *Gpnmb* upregulation by MITF^90–92^ may represent a critical node in the crosstalk between lysosomes and the inflammatory tone of myeloid cells. Because GPNMB is highly upregulated in response to lysosomal perturbations and is necessary for proper lysosomal acidification, we speculate that it functions not only as an anti-inflammatory signaling molecule^93^, but also as an intracellular lysosomal pH-responsive rheostat, engaging mechanisms to restore lysosomal ion homeostasis to further dampen immune responses arising from lysosomal dysfunction.

During normal development and aging, macrophages and microglia engulf many extracellular substrates for clearance, which ultimately requires sustained lysosomal function. This demand is particularly high in the context of many neurodegenerative diseases, in which the high cargo burden can eventually exceed the lysosome’s degradative capacity, resulting in the accumulation of aberrant protein and lipid species that are thought to contribute to disease pathogenesis and progression. Because the dynamic regulation of lysosomal pH is a critical determinant of the capacity of myeloid cells to clear engulfed cargo, such as β-amyloid^94–96^, alterations in the ability to properly control lysosomal pH may not only drive myeloid cell state changes downstream of lysosomes, but also secondarily by contributing to the availability of pathological species competent in activating previously reported signaling pathways. Although the microglial subpopulation that emerges in response to lysosomal deacidification expresses numerous unique marker genes that distinguish it from previously reported microglial states in other models, it is curious that many genes that are concomitantly induced as part of this response have been similarly identified as part of the DAM signature in other contexts, including in AD mouse models and human post-mortem brain tissue^10,97,98^. Similarly, the TFs nominated to regulate the microglial response in models of severe lysosomal dysfunction (this manuscript and Balak *et al.*) were highly overlapping with those predicted to mediate the responses to amyloid and tau/APOE4 (accompanying paper, Schlachetzki *et al.*), suggesting potential converging cellular mechanisms could be involved. It is possible that, in addition to receptor-dependent signaling mechanisms previously described, an age- and disease-dependent decline in microglial lysosomal health in these contexts may also contribute to the observed microglial states; however, a comprehensive assessment of lysosomal properties in microglia from these models is required. Because lysosomal deficits are prevalent in many neurodegenerative diseases, either due to cell autonomous deficiencies resulting from genetic mutations, or secondarily through age-associated impairments of lysosomal function, it is likely that the findings we report here are relevant to many other neurodegenerative conditions as well.

## METHODS

### Animal husbandry

Wild-type (WT) mice were obtained from Jackson Laboratories or Charles River Laboratories for primary cell derivation. For mouse bone marrow derived macrophages (BMDMs), 12 week old male mice on a C57Bl/6J background (Jackson Laboratories; Cat# 000664) were used. For primary mouse microglia, timed pregnant E14 female mice on a C57Bl/6NCrl background (Charles River Laboratories; Cat# 027) were used. For animals used for single-cell sequencing studies, *Grn^+/+^* and Grn^-/-^ (*Grn* KO) mice (on a C57Bl/6J background with hTfR^mu/hu^ knock-in)^42^ were aged to 24 months. Mice were housed under a 12-hour light/dark cycle and group housed. Mice were housed with enrichment and bedding was changed weekly. Animals were monitored weekly for health/immune issues before and during studies. And excluded if serious concerns were observed. All mouse procedures adhered to regulations and protocols approved by Denali Therapeutics Institutional Animal Care and Use Committee.

### Sorting of fresh microglia of aged mice for scRNA-seq

Two year old mice were anesthetized, transcardially perfused with PBS, and brains were removed for scRNA-seq or microglial H3K27ac analyses (see below). Hemibrains were separated and one half was frozen for nuclei isolation for H3K27ac ChIP-seq, and the other was diced into a fine slurry and dissociated using the Miltenyi Adult Brain Dissociation Kit (Miltenyi Biotec, Cat# 130- 107-677) in the presence of transcription and translation inhibitors 5μg/ml actinomycin D (Cell Signaling Technology; Cat# 15021) and 0.56μg/ml anisomycin (Selleck Chemical; Cat# S7409). After dissociation and debris removal, cells were stained on ice for 30 minutes, protected from light. The following primary antibodies were used: anti-CD11b BV421 (BD; Cat# 562605; 1:100), anti-CD45 APC (Biolegend; Cat# 157606; 1:100), anti-CD44 PE-Cy7 (Sigma-Aldrich; MABF426; 1:100), anti-CD163 PE-Cy7 (BioLegend; Cat# 155320; 1:100), and anti-Ly6G PE-Cy7 (BioLegend; Cat# 127618; 1:100). Total live cell counts were measured using CountBright Plus Absolute Counting Beads (Invitrogen; Cat# C36995) by flow using a FACS Aria III (BD). Equal numbers of cells from all samples were pelleted and labeled using CellPlex CMO tags (10x Genomics, CG000391 Rev A) for multiplexing, following manufacturer protocols. Finally, an equal number of cells per sample were combined into one pool for FACS sorting and sorted and CD11b^+^/CD45^+^/CD44^-^/CD163^-^/Ly6G^-^ cells were collected, pelleted, counted, and processed for scRNA-seq.

### 10x Genomics scRNA-seq and analysis

The pool of FACS sorted cells was loaded into 2 replicate GEM channels on the microfluidic Chromium Next GEM Chip G for single cell capture per manufacturer protocol (10X Genomics, CG000388_Rev B). 50,000 cells were loaded per GEM channel for a target capture of 30,000 cells each, or a combined target capture of 4,000-6,000 cells per animal. After cell capture, RT, and cDNA amplification, each replicate was separated into one gene expression (GEX) library and one sample tag (CMO) library via dual-sided SPRI bead purification. Library preparation was performed per manufacturer’s protocol and libraries for both replicates were combined equimolar. Sequencing was performed by SeqMatic (Fremont) on 2 lanes of an Illumina NovaSeq X 10B cartridge (28X10X10X90), targeting 50,000 reads per cell for mRNA transcripts and 5,000 reads per cell for demultiplexing of CellPlex oligos.

Raw scRNA-seq reads were processed with Cell Ranger v7.0.1. Two pooled libraries were retained, and samples were demultiplexed by CMO distribution using the “cellranger multi” function. Filtered expression matrices were further analyzed in R v4.3. Analysis was performed primarily using Seurat v5^99^. Droplets with fewer than 200 unique genes and genes found in fewer than 3 droplets were discarded. To remove batch effects, CCA integration was performed between the 2 pools of samples. Normalization and dimensionality reduction were carried out according to the standard Seurat v5 pipeline. Putative doublet droplets were identified and removed using the “scDblFinder” package. Major cell types were determined by clustering and marker gene analysis. Microglia were isolated and subclustered for further analysis. For all downstream analysis, 9000 cells were randomly selected from each genotype to eliminate capture biases in population distributions. Microglia clusters were characterized by examining marker genes calculated by Wilcoxon Rank Sum test. Differences in cluster distribution across experimental groups was determined using the “propeller” function in the “speckle” package^100^, which applies a logit transformation and determines statistical significance of proportional shifts.

### Immunohistochemistry, slide scanner, and super-resolution microscopy of mouse tissue

Mouse brains were prepared for immunohistochemistry as previously described^42^. Briefly, mice were anesthetized and transcardially perfused with PBS, followed by 4% paraformaldehyde (Electron Microscopy Sciences; Cat# 15714-S). Mouse hemibrains were sectioned coronally using Multibrain sectioning service (Neuroscience Associates) at a thickness of 40μm. Gelatin sheets containing embedded sections (40 animals/sheet) were stored in antigen preservation solution (50% PBS: 50% ethylene glycol, 1% PVP) until staining. Sheets containing sections were washed three times in cold PBS for 5 minutes each, then blocked in universal blocking buffer (1% BSA, 1x fish gelatin, 0.5% Triton X-100, 0.1% sodium azide) for 1 hour at room temperature with gentle agitation. After blocking, sections were incubated in primary antibody diluted in antibody dilution buffer (1% BSA in PBS, 0.3% Triton X-100, 0.1% sodium azide) overnight at 4°C. The following day, sections were washed three times in PBST for 10 minutes each. Next, sections were incubated in secondary antibody diluted in antibody dilution buffer at room temperature for one hour, protected from light. Sections were then washed in PBST containing DAPI for 10 minutes, then two more times in PBST for 10 minutes each. Finally, sheets containing brain sections were mounted onto slides using ProLong Glass Antifade Mountant (Invitrogen; Cat# P36984). The following primary antibodies were used: rabbit anti-GPNMB (Cell Signaling Technology; Cat# 90205; 1:500) and guinea pig anti-IBA1 (SYSY; Cat# 234308; 1:1000). The following secondary antibodies were used: donkey anti-guinea pig IgG AlexaFluor 647 (Jackson Immunoresearch; Cat# 706-605-148; 1:500), donkey anti-rabbit IgG AlexaFluor 750 (Abcam; Cat# ab175731; 1:500), and donkey anti-rabbit IgG AlexaFluor 594 (Invitrogen; Cat# A-21208; 1:500).

For quantification, slides were imaged using a Zeiss Axio Scan.Z1 digital slide scanner (Zeiss) running Zen 3.7 software (Zeiss) using a 20x or 40x objective. Each tissue section was entirely imaged by scanning across the identified tissue area, taking images of each location with exposure settings and filter sets to detect DAPI, autofluorescence (488), autofluorescence (594), Alexa-647-, and Alexa-750-tagged antibodies. Automated shading correction and stitching in Zen was then used to produce single complete images for each section. Identical imaging settings and post-processing methods were used to collect images from all tissue sections from the experiment in a single imaging run. Standardized anatomical locations within the tissue were manually outlined in the collected images, avoiding tissue fold or mounting media bubble artifacts, in the generated images. To quantify IBA1, GPNMB, and lipofuscin (autofluorescence) localization and distribution across the selected areas in the tissue sections, images were first processed to identify specific classes of objects using sequences of advanced image processing tools in custom macros designed in-house with Zen 3.7 software (Zeiss). GPNMB, IBA1, and lipofuscin (both 594 and 488) stained objects were detected by local thresholding and size selection to remove too-small staining artifacts; objects smaller than cell body and process-sized objects removed for IBA1, smaller than cell body sized objects removed for all other channels. Colocalization objects were then defined as areas of mutual overlap between two or more sets of single-channel identified objects. IBA1-nonGPNMB objects were defined as areas of IBA1 objects that did not colocalize with GPNMB objects. The total area (μm^2^) of the tissue analyzed was defined by the areas manually selected and outlined. Only single-channel and overlap objects within areas of interest were included in the analyses; cortex, hippocampus, and thalamus areas were separately quantified in each tissue section. Once classes of objects (single-channel and overlap) were identified, we quantified the total count, total area, and total sum intensity of the Alexa750 (GPNMB) channel for the total tissue area and each set of objects in every area selected within each imaged tissue section. Total sum intensity of Alexa488 (lipofuscin) was also quantified for a subset of objects.

For super-resolution imaging, fixed and stained brain sections were mounted on the slide and imaged on a Leica SP8 microscope using a 63x/1.4 N.A. oil immersion objective with a pixel size of ∼30nm at 63x magnification in lightning acquisition mode. For acquisition of super - resolution images, lightning processing was performed in the adaptive mode and images were then visualized and processed in LAS X and/or ImageJ.

### Sorting of fresh microglial nuclei of aged mice for ChIP-seq

Brain nuclei were isolated as previously described^101,102^, with snap-frozen hemibrains from WT and *Grn* KO mice homogenized in 1% formaldehyde in Dulbecco’s phosphate buffered saline. Nuclei were stained overnight at 4°C with PU.1-PE conjugated antibody (Cell Signaling Technology; Cat# 81886S). Following antibody incubation, nuclei were washed with 4mL FANS buffer, passed through a 40μm just prior to sorting. Nuclei were sorted with a BD Influx Cell Sorter gating for PU.1^+^/NeuN^-^/OLIG2^-^ nuclei. Following sorting, nuclei were pelleted at 1,600xg for 5 minutes at 4°C in FANS buffer, supernatant was aspirated to 500μL, then resuspended and transferred to 1.5mL tubes. These were spun at 1,600xg for 15 minutes at 4°C and all supernatant removed. Nuclei pellets were snap frozen and stored at -80°C prior to library preparation.

### H3K27ac ChIP-seq and analysis

Chromatin immunoprecipitation was performed as previously described^103,104^. Briefly, ∼500,000 fixed nuclei were quickly thawed on ice and resuspended in ice-cold LB3 buffer (10mM Tris-HCl pH 7.5, 100mM NaCl, 1mM EDTA, 0.5mM EGTA, 0.1% Na-Deoxycholate, 0.5% N-lauroylsarcosine) and 1x Protease Inhibitor Cocktail (Sigma). Nuclei/chromatin were sheared by sonication using the PIXUL Multi-Sample Sonicator (Active Motif) in low bind 96 well plates (Corning; Cat# 7007) following the manufacturer’s instructions. Samples were sonicated with the following settings: Pulse (N): 50, PRF [kHz]: 1.00, Process time: 60 min, Burst Rate [Hz]: 20.00. Samples were pulse spun to 600xg at 4°C in the 96-well plate, transferred to Eppendorf tubes, and chromatin was recovered by retaining the supernatant following a spin at 21,000xg at 4°C for 10 minutes. The supernatant was then diluted 1.1-fold with ice-cold 10% Triton X-100. Two percent of the lysate was kept as ChIP input control. 10μL of Dynabead Protein A and 10μL of Protein G were added per sample, in addition to 2μg of an antibody specific for H3K27ac (Active Motif; Cat# 39685). The samples were rotated overnight at 4°C and were washed as follows the next day: 3x with Wash Buffer I (20mM Tris-HCl pH 7.5, 150mM NaCl, 2mM EDTA, 0.1% SDS, 1% Triton X-100) + protease inhibitor cocktail, 3x with Wash Buffer III (10mM Tris-HCl pH 7.5, 250mM LiCl, 1% Triton X-100, 1mM EDTA, 0.7% Sodium Deoxycholate) + protease inhibitor cocktail, 2x with TET (0.2% Tween-20/TE) + 1/3 protease inhibitor cocktail, and 1x with IDTET (0.2% Tween-20, 10mM Tris pH8, 0.1mM EDTA). Samples were finally resuspended in 25μL TT buffer (10mM Tris pH 8 + 0.05% Tween 20) prior to on-bead library preparation. Libraries were prepared with the NEBNext Ultra II DNA library prep kit (NEB) reagents according to the manufacturer’s protocol on the beads suspended in 25μL TT buffer, with reagent volumes reduced by half. DNA was eluted and crosslinks reversed by adding 4μL 10% SDS, 4.5μL 5 M NaCl, 3μL EDTA, 4μL EGTA, 1μL proteinase K (20mg/ml), 16μL water, incubating for 1 h at 55°C, then overnight at 65°C. DNA was purified using 2μL of SpeedBeads (GE Healthcare) in a 20% PEG8000 / 1.5 M NaCl solution to final concentration of 12% PEG, then eluted with 25μl TT. DNA contained in the eluate was then amplified for 14 cycles in 25μl PCR reactions using NEBNext High-Fidelity Q5 2X PCR Master Mix (NEB) and 0.5 mM each of Solexa 1GA and Solexa 1GB primers. The resulting libraries were size selected by gel excision to 200-500 bp, purified, and paired-end sequenced (SP100) to a depth of ∼20 million reads using an Illumina NovaSeq instrument.

FASTQ files from ChIP sequencing experiments were mapped to the mouse mm10 genome using Bowtie2 with default parameters^105^. HOMER was used to convert reads aligning to a single genomic location into “tag directories” for subsequent analysis^106^. HOMER was also used to generate sequencing-depth normalized bedGraph/bigWig files for visualization in the UCSC Genome Browser (http://genome.ucsc.edu). To quantify differential regions of H3K27 acetylation between genotypes, an ATAC peak file was created with HOMER’s mergePeaks command using published data from Fixsen et al.^107^ and the accompanying manuscript (Balak *et al*.) to represent a broad range of open chromatin regions in microglia across several conditions. These ATAC-seq peaks were annotated with raw tag counts with HOMER’s annotatePeaks using parameters -noadj and -size 800. Subsequently, DESeq2^108^ was used to identify the differential H3K27ac signal with the cutoffs of FC>2 and p-adj<0.05. To identify motifs enriched in differential peak regions over background, HOMER’s motif analysis (findMotifsGenome.pl) including known default motifs and *de novo* motifs was used. Background sequences were comprised of random GC-matched genomic sequences.

### Isolation of primary mouse microglia

For isolation of primary microglia, brains of neonatal mice (P1-P3) were removed and the cerebral cortices and hippocampus were harvested. Isolated tissues were diced into a fine slurry, suspended in 50mL cold HBSS, and centrifuged at 200xg for 3 minutes. Then, the pelleted tissue was resuspended in warm 0.25% Trypsin-EDTA (Gibco; Cat# 25200056), mixed by inversion, and incubated at 37°C for 20 minutes with agitation (100rpm). After 20 minutes, the enzymatic digestion was stopped by adding primary microglia culture medium [DMEM/F12 (Gibco; Cat# 11- 330-057), 10% HyClone FBS (Cytiva; Cat# SH30080.03), 1X GlutaMax (Gibco; Cat# 35050061), and penicillin-streptomycin (Gibco; Cat# 15140122)] supplemented with DNase-I (Sigma-Aldrich; Cat# 5025) at 625U/mL culture medium. Dissociated tissue was mixed by inversion 10 -12 times and triturated until a homogenous, single-cell suspension was observed. Cells were pelleted by centrifugation at 200xg for 7 minutes, supernatant was removed, and the cells were resuspended in primary microglia culture medium. Cells were filtered through a 70μm strainer to create a single-cell suspension and counted. Poly-D-lysine (Sigma-Aldrich; Cat# P6407)-coated T-75 vented culture flasks were washed twice with PBS and primary microglia were seeded at a density of 6 million cells/flask.

After 3 days, the medium was replaced with fresh primary microglia culture medium. Subsequently, half-media changes were performed every other day until the mixed glial culture reached a confluency of 90% (10-14 days). Microglia were then detached from the mixed glial flask using a heated orbital shaker for 2 hours at 240rpm at 37°C and 5% CO_2_. The microglial suspension was centrifuged at 200xg for 8 minutes, resuspended in conditioned medium, counted, and seeded for experiments in 50% conditioned medium with 50% fresh medium. The original mixed glial culture flasks were replenished with fresh medium and maintained in culture with half-media changes every other day. Primary microglia were similarly harvested every 6-8 days for a total duration of 3 weeks.

Isolated primary microglia were seeded at assay-dependent concentrations onto poly-D-lysine-coated plates in 50% conditioned medium from the mixed glial culture flask with 50% fresh primary microglia culture medium [DMEM/F12 (Gibco; Cat# 11-330-057), 10% HyClone FBS (Cytiva; Cat# SH30080.03), 1X GlutaMax (Gibco; Cat# 35050061), and penicillin-streptomycin (Gibco; Cat# 15140122)]. After 24 hours, seeded microglia were washed twice with PBS to remove debris, cultured in 50% fresh medium and 50% conditioned medium (collected from mixed glial culture flasks and centrifuged at 1000xg for 3 minutes to remove cellular debris), and used for experiments. For experiments involving pharmacological treatments, DMSO-based solutions were dispensed using a Tecan D300e dispenser (Tecan), and all other solutions were dosed manually.

### Derivation of primary mouse BMDMs

Primary mouse BMDMs were prepared as described previously^42,64^ from 12 week old WT or *Grn^-/-^* animals, with slight modifications. Briefly, animals were sacrificed and the femur and tibia bones were collected into cold HBSS (Gibco; Cat# 14175095). Bones from multiple mice (6 for WT only studies, 3 for WT/*Grn* KO mice) were pooled for each individual biological replicate and sterilized with 70% isopropanol, washed three times with HBSS, and cracked in HBSS using a mortar and pestle. The cell suspension containing bone marrow cells was filtered through a 70μm strainer to remove cracked bone pieces and centrifuged at 300xg for 5 minutes. The supernatant was removed, and the pellet was resuspended in ACK Lysing Buffer (ThermoFisher; Cat# A1049201) for 2 minutes at room temperature to remove red blood cells. Then, BMDM culture medium [RPMI- 1640 (Gibco; Cat# 61870036), 10% HyClone FBS (Cytiva; Cat# SH30080.03), and penicillin - streptomycin (Gibco; Cat# 15140122)] was added to stop the ACK lysis reaction. Cells were pelleted at 300xg for 5 minutes and the supernatant was discarded. Then, the cell pellet was resuspended in BMDM medium, counted, and plated onto 15cm non-tissue culture treated petri dishes at a density of 20 million cells/plate in BMDM medium containing murine M-CSF (Life Technologies; Cat# PMC2044) at 50ng/mL. Four days after seeding the cells, an additional 50ng/mL fresh M-CSF was added to each plate. On day 7, differentiated BMDMs were washed once with PBS and harvested in BMDM medium using a cell scraper. Cells were pelleted at 300xg for 5 minutes, resuspended in BMDM medium, and counted. Finally, cells were centrifuged at 300xg for 5 minutes, resuspended in Recovery Cell Culture Freezing Medium (Gibco; Cat# 12648010) at a density of 10 million cells/mL, and cryopreserved for later use.

Differentiated BMDMs were thawed, counted, and seeded at assay-dependent concentrations onto poly-D-lysine-coated plates in BMDM culture medium [RPMI-1640 (Gibco; Cat# 61870036), 10% HyClone FBS (Cytiva; Cat# SH30080.03), and penicillin-streptomycin (Gibco; Cat# 15140122)] with supplemental M-CSF (50ng/mL), unless otherwise noted. Fresh BMDM medium was changed 24 hours after seeding of the cells and utilized for experiments, as detailed below. For experiments involving pharmacological treatments, DMSO-based solutions were dispensed using a Tecan D300e dispenser (Tecan), and all other solutions were dosed manually. All compounds used were confirmed to be endotoxin free (<0.1 EU/mL).

### RNA sample and library preparation for bulk RNA-sequencing

RNA was harvested from cells using the RNeasy Micro Kit (Qiagen), following the manufacturer’s protocol. RNA integrity (RIN) and concentration were measured by High Sensitivity RNA Screen Tape Analysis (Agilent) on a 4200 TapeStation System (Agilent). Libraries were generated using the QuantSeq 3’ mRNA-seq Library Prep Kit FWB for Illumina (Lexogen A01173) with the UMI second strand synthesis module to identify and remove PCR duplicates, following the protocol defined by the manufacturer. Library quantity and quality were assessed with TapeStation High Sensitivity D1000 Screen Tape (Agilent 5067-5592) using High Sensitivity D1000 reagents (Agilent 5067-5593). Libraries were pooled in equimolar ratios and sent to SeqMatic for sequencing on a NovaSeq S1 (75bp, single-end reads).

### Bulk RNA-seq analysis

Raw bulk RNA-seq sequencing data was processed using nf-core/rnaseq v3.11.2 (doi: https://doi.org/10.5281/zenodo.1400710) of the nf-core collection of workflows^109^, executed with Nextflow v23.10.0^110^. The M31 Gencode release of the GRCm39 mouse genome was used as the reference to map reads and quantify gene-level expression. Because the QuantSeq-UMI library preparation captures 3’ tags of transcripts, the following custom arguments were passed to the STAR alignment step: ’--alignIntronMax 1000000 --alignIntronMin 20 --alignMatesGapMax 1000000 --alignSJoverhangMin 8 --outFilterMismatchNmax 999 --outFilterType BySJout -- outFilterMismatchNoverLmax 0.1 --clip3pAdapterSeq AAAAAAAA’ and the salmon quantification was performed without length correction.

Downstream analysis of the count matrices returned by the nf-core/rnaseq workflow was performed using R version 4.3.1. The edgeR::normLibSizes function was used to calculate library sizes^111^ using the TMM method^112^. Afterwards, the data was subset to protein-coding genes and a linear model was fit with the edgeR::voomLmFit function, specifying the experimental group, sample collection batch (if any) and differences in library preparation protocol (if any) as main effects. Differential expression results were obtained with the limma::makeContrasts, limma::contrasts.fit and limma::eBayes functions, the latter was robustified against outliers with the robust=TRUE argument^113^.

### Protein extraction and western blotting

For immunoblotting from cultured cells, medium was first removed and RIPA buffer (Teknova ; Cat# R3792) supplemented with cOmplete, Mini, EDTA-free protease inhibitor cocktail (Sigma-Aldrich; Cat# 4693159001) and PhosSTOP phosphatase inhibitor (Sigma-Aldrich; Cat# 4906845001) was added to each well. Cell lysates were scraped and collected, followed by rotation for 15 minutes at 4°C. Cell debris was removed by centrifugation at 21,000xg for 10 minutes at 4°C. The supernatant was transferred to Protein LoBind tubes (Eppendorf; Cat# 13- 698-794) and subjected to BCA analysis for total protein concentration determination using the Pierce BCA Protein Assay Kit (Thermo Scientific; Cat# 23225). Equal total protein lysates were diluted in NuPAGE Sample Reducing Agent (Invitrogen; Cat# NP0009) and NuPage LDS Sample Buffer (Invitrogen; Cat# NP0007) and boiled at 95°C for 10 minutes. Denatured protein lysates were cooled to room temperature and used for downstream immunoblotting assays.

For all immunoblotting experiments, 20μg total protein lysate was loaded per sample onto NuPAGE 4 to 12%, Bis-Tris, 1.0mm protein gels (Invitrogen; Cat# WG1402, WG1403) in NuPAGE MOPS SDS Running Buffer (Invitrogen; Cat# NP0001) or MES buffer (Invitrogen; Cat# NP0002). Proteins were then transferred using Trans-Blot Turbo Midi 0.2μM Nitrocellulose Transfer Packs (Bio-Rad; Cat# 1704159) with a Trans-Blot Turbo Transfer System (Bio-Rad), or using with iBlot3 Midi Nitrocellulose Transfer Packs (ThermoFisher; Cat# IB31001) with an iBlot3 Western Blot Transfer System (ThermoFisher). Membranes were washed three times with TBS-T for 5 minutes each on a rocking platform, blocked in Blocking Buffer for Fluorescent Western Blotting (Rockland; Cat# MB-070) for 1 hour at room temperature on a rocking platform, and incubated with primary antibodies diluted in Rockland Blocking Buffer overnight at 4°C on a rocking platform. The following day, membranes were washed three times with TBS-T for 5 minutes each on a rocking platform, incubated in secondary antibodies diluted in Rockland Blocking Buffer at room temperature on a rocking platform protected from light, washed three times with TBS-T for 5 minutes each on a rocking platform protected from light, and visualized using an Odyssey CLx Imager (LI-COR Biosciences). Immunoblots were analyzed using ImageStudio (LI-COR Biosciences).

The following primary antibodies were used: sheep anti-PGRN (R&D Systems; Cat# 2557; 0.5μg/mL), rabbit anti-GPNMB (Cell Signaling Technology; Cat# 90205; 1:500), rabbit anti-FABP5 (Cell Signaling Technology; Cat# 39926; 1:500), rabbit anti-CatB (Cell Signaling Technology; Cat# 31718; 1:500), rabbit anti-LPL (Biotechne; Cat# AF7197; 1μg/mL), mouse anti-Vinculin (Sigma-Aldrich; Cat# V9264; 1:1000), mouse anti-β-Actin (Sigma-Aldrich; Cat# A2228; 1:10000), mouse anti-LC3 (Cell Signaling Technology; Cat# 83506; 1:500), mouse anti-p62 (Biotechne; Cat# MAB8028; 2μg/mL), mouse anti-AMPKα (Cell Signaling Technology; Cat# 2793; 1:500), rabbit anti-phospho-AMPKα (T172) (Cell Signaling Technology; Cat# 2535; 1:500), mouse anti-GAPDH (Abcam; Cat# ab8245; 1:2000), rabbit anti-ATP6V1A (Cell Signaling Technology; Cat# 39517; 1:500), rabbit anti-NPC1 (Abcam; Cat# ab134113; 1:500), rabbit anti-VPS34 (Cell Signaling Technology; Cat# 4263; 1:500), rabbit anti-CatD (Cell Signaling Technology; Cat# 88239; 1:500), sheep anti-TREM2 (Biotechne; Cat# AF1729; 0.1μg/mL), rabbit anti-ATG7 (Cell Signaling Technology; Cat# 8558; 1:500), rabbit anti-ATG14 (Cell Signaling Technology; Cat# 96752; 1:500), and rabbit anti-UVRAG (Abcam; Cat# ab313627; 1:500). The following secondary antibodies were used: IRDye 680RD goat anti-mouse IgG (LI-COR; Cat# 926-68070; 1:15000), IRDye 680RD goat anti-rabbit IgG (LI-COR; Cat# 926-68071; 1:15000), IRDye 800CW goat anti-mouse IgG (LI-COR; Cat# 926-32210; 1:15000), IRDye 800CW goat anti-rabbit IgG (LI-COR; Cat# 926-32211; 1:15000), and rabbit anti-sheep IgG DyLight800 (Invitrogen; Cat# SA5-10060; 1:15000).

### Measurement of lysosomal pH

Lysosomal pH was measured by a pulse-chase and ratiometric fluorescence measurement of OregonGreen-conjugated dextran loaded into cells via fluid-phase endocytosis and imaged at pH-sensitive (signal) and pH-insensitive (loading control) wavelengths. Briefly, fresh medium containing 50μg/mL 10 kDa dextran-OregonGreen (OG) (Invitrogen; Cat# D7170) was added onto cells. The next day, media was changed to remove excess extracellular dextrans. After 110 minutes, medium on selected wells for pH standard calibration was removed and replaced with calibration buffers (10mM HEPES, 10mM MES, 10mM sodium acetate, 140mM KCl, 5mM NaCl, 1mM MgCl_2_) ranging from pH 3.75 to 7.5 and containing 30μM nigericin (Invivogen; Cat# tlr-nig) and 15μM monensin (Sigma-Aldrich; Cat# M5273) ionophores, or with test solution (calibration solution at physiological pH without supplemental ionophores). After another 10 minutes (total chase period of 120 minutes), live imaging was performed on the calibration and test wells using a PerkinElmer Opera Phenix high content microscope at 40X magnification, imaging OG at two wavelengths (Exc 488nm, Em 500-550nm; pH-sensitive) and (Exc 425nm, Em 435-550nm; pH-insensitive). A standard curve for lysosomal pH was generated using mean fluorescence intensity of 488/425, which was used to calculate pH for test wells. Mean fluorescence intensity per cell was quantified over 13 FOVs per condition per biological replicate.

### Measurement of GCase activity

Lysosomal GCase activity was measured using a LysoFQ-GBA activity probe^114^. Briefly, cells were plated at a density of 75,000 cells/well. On the day of the experiment, BMDM media was changed and control wells were treated with 100μM GCase inhibitior conduritol B epoxide (CBE; Sigma-Aldrich; Cat# C5424). After a one hour pre-treatment with CBE, all wells were spiked with LysoFQ-GBA at a final concentration of 5μM. Wells were immediately imaged and mean fluorescence intensity per cell was quantified over 6 FOVs per condition per biological replicate.

### Immunocytochemistry

For all high-content imaging experiments, primary BMDMs or microglia were plated at a density of 75,000 cells/well. Unless otherwise specified, cells were washed twice with cold PBS and fixed in 4% PFA (Electron Microscopy Sciences; Cat #15714) diluted in PBS. Cells were then washed twice with cold PBS, permeabilized with permeabilization/wash buffer [0.05% saponin (Sigma-Aldrich; Cat# SAE0073) diluted in PBS] for 20 minutes, and blocked with blocking buffer [5% bovine serum albumin (BSA) (Sigma-Aldrich; Cat# A8412) diluted in permeabilization/wash buffer] for 1 hour. Cells were then incubated in primary antibodies diluted in blocking buffer overnight at 4°C on a rocking platform. On the next day, wells were washed twice with permeabilization/wash buffer and incubated in secondary antibodies diluted in blocking buffer for 1 hour at 25°C on a rocking platform, protected from light. Next, wells were washed twice with permeabilization/wash buffer, and cells were incubated in 1mM DAPI in PBS for 10 minutes, protected from light. Finally, cells were washed twice with PBS and imaged using a PerkinElmer Opera Phenix high content microscope. The following primary antibodies were used: mouse anti-Galectin-3 (ab2785; Cat# ab2785; 1:100), rabbit anti-CHMP4B (Proteintech; Cat# 13683-1-AP; 1:200), and rat anti-LAMP2 (abcam; Cat# ab13524; 1:250). The following secondary antibodies were used: goat anti-rat IgG 488 (Invitrogen; Cat# A-11006; 1:300), goat anti-rabbit IgG 594 (Invitrogen; Cat# A-11037; 1:300), goat anti-mouse IgG 647 (Invitrogen; Cat# A-21235; 1:300). High-content imaging was performed using a PerkinElmer Opera Phenix high content microscope and analyzed using Harmony High Content Analysis Software (PerkinElmer). For super-resolution imaging, fixed and stained cells in 96-well plates were rinsed twice with PBS prior to imaging on a Leica SP8 microscope using a 63x/1.4 N.A. oil immersion objective with a pixel size of ∼30nm at 63x magnification in lightning acquisition mode. For acquisition of super-resolution images, lightning processing was performed in the adaptive mode and images were then visualized and processed in LAS X and/or ImageJ.

### Measurement of lysosomal proteolysis

Lysosomal proteolysis was measured using several approaches. For DQ-BSA assays, fresh BMDM medium (without supplemental M-CSF) containing NucBlue Live ReadyProbes and lysosomal drugs was changed 24 hours after cell plating. Cells were pre-treated with lysosomal drugs for 22 hours. After 22 hours, fresh BMDM medium containing NucBlue Live ReadyProbes and 10μg/mL DQ green BSA (Invitrogen; Cat# D12050) and 10μg/mL BSA-AlexaFluor647 (Invitrogen; Cat# A34785) were added to each well. Lysosomal drugs were immediately added to each respective well and plates were imaged live using a PerkinElmer Opera Phenix high content microscope at 63X magnification. Images were acquired every 30 minutes for 13 hours. Mean DQ-BSA intensity relative to BSA-647 per cell was quantified over 12 FOVs per condition per biological replicate.

Proteolytic activity of specific lysosomal cathepsins was also probed. For cathepsin activity assays, fresh BMDM medium (without supplemental M-CSF) and lysosomal drugs was changed 24 hours after cell plating. Cells were treated with lysosomal drugs for 22 hours. After 22 hours, NucBlue Live Ready Probes and 1X Green Cathepsin B Assay (Immunochemistry Technologies; Cat# 9151) was spiked into each sample well and incubated for 30 minutes. Cells were then washed twice with PBS and imaged live immediately in Live Cell Imaging Solution (Invitrogen; Cat# A14291DJ) using a PerkinElmer Opera Phenix high content microscope at 63X magnification. Mean Green Cathepsin B or spot intensity per cell was quantified over 31 FOVs per condition per biological replicate.

### Cytokine analysis

For analysis of secreted cytokines and chemokines from cultured cells, medium containing lysosomal drugs was changed 24 hours after cell plating. After 21 hours of treatment, NucBlue Live ReadyProbes were spiked in to each well, incubated for 30 minutes, and imaged live using a PerkinElmer Opera Phenix high content microscope at 10X magnification to determine nuclei number. Nuclei count was quantified over 9 FOVs per condition per biological replicate. After measuring nuclei number, supernatants were harvested into Protein LoBind Tubes (Eppendorf; Cat# 0030108434), flash frozen, and exported for processing by Eve Technologies (Canada) using their Mouse Cytokine/Chemokine 44-plex Discovery Assay Array (Eve Technologies; Cat# MD44) in duplicate. For CRISPR knockout experiments, media was conditioned between days 11-13 post-transfection, nuclei were counted, and supernatant was harvested for cytokine/chemokine analysis. Cytokine concentrations from duplicate runs for averaged for each sample and were normalized to total nuclei count for the well from which the supernatant was harvested from.

### Pharmacological treatments for in vitro experiments

For all experiments involving pharmacological treatment of primary mouse BMDMs or microglia, a fresh medium change was performed 24 hours after plating of the cells. Unless otherwise specified, drugs were added using a digital dispenser (Tecan D300e) for DMSO-based compounds or manually (non-DMSO-based compounds). The following compounds were used: conduritol B epoxide (CBE) (Sigma-Aldrich; Cat# C5424), Bafilomycin A1 (BafA1) (Cell Signaling Technology; Cat# 54645), U18666A (Bio-Techne; Cat# 1638), VPS34IN1 (Selleck Chemical; Cat# S7980), E64 (Selleck Chemical; Cat# 50-136-4719), leupeptin hemisulfate (Sigma-Aldrich; Cat# 108-976), pepstatin A (Enzo Life Sciences; NC1460915), monosodium urate (MSU) crystals (Invivogen; Cat# NC0872699), and rimeporide (Selleck Chemicals; Cat# 50-234-1204).

### Measurement of free cholesterol

Cholesterol was measured using a commercial cholesterol assay kit (Abcam; Cat# ab133116), according to manufacturer’s instructions. After removing culture medium from sample wells, cells were fixed for 10 minutes with the Cell-Based Assay Fixative solution. Cells were then washed three times with Cholesterol Detection Wash Buffer. Filipin III (dissolved in ethanol) was then diluted 1:100 in Cholesterol Detection Assay Buffer, NucRed Live 647 ReadyProbes (Invitrogen; Cat# R37106) were added, and 100mL total volume was added to each well, followed by incubation for 60 minutes protected from light. Then, cells were washed twice with Cholesterol Detection Wash Buffer, and imaged immediately using a PerkinElmer Opera Phenix high content microscope at 63X magnification. Mean number of cells containing membrane or cytosolic filipin III-positive spots was quantified over 13 FOVs per condition per biological replicate.

### Analysis of endolysosomal vesicle size and vacuolation

Endolysosomal vesicle size was determined using fluorescently-labeled dextrans. Briefly, fresh culture medium containing 33μg/mL 3 kDa dextran-AlexaFluor 488 (Invitrogen; D34682), 33 μg/mL 10 kDa dextran-AlexaFluor 647 (Invitrogen; D22914), and 33μg/mL 70 kDa dextran-tetramethylrhodamine (TMR) (Invitrogen; Cat# D1818) was added onto cells 24 hours after plating. The following day, fresh culture medium containing NucBlue LiveReady Probes and various pharmacological agents was changed. After a chase/treatment period of 120 minutes, cells were imaged live using a PerkinElmer Opera Phenix high content microscope at 63X magnification. Mean spot size for each labeled dextran (marking endolysosomes that were present prior to treatment with different drugs) was quantified over 81 FOVs (each FOV containing ∼21 cells on average) per condition per biological replicate.

### EdU incorporation assay

EdU incorporation assays were performed using an EdU Staining Proliferation Kit (Abcam; ab219801) according to manufacturer’s instructions, with slight modifications. Fresh BMDM medium containing 1μM EdU was added to cells 24 hours after plating immediately prior to the addition of various lysosomal drugs. After treatment with lysosomal drugs for 22 hours in the presence of EdU, cells were washed twice with cold PBS and fixed using 4% paraformaldehyde (PFA) (Electron Microscopy Sciences; Cat #15714) in PBS for 15 minutes at room temperature. Cells were then washed twice with wash buffer (3% BSA in PBS) and permeabilized using 1X Permeabilization buffer for 20 minutes at room temperature. After permeabilization, cells were washed twice with wash buffer. Then, 1X reaction mix (4mM copper sulfate, 1.25μM iFluor 488 azide, 2μg/mL sodium ascorbate in PBS) was added to each well and cells were incubated for 1 hour at 25°C on a rocking platform, protected from light. Finally, cells were washed twice with wash buffer prior to proceeding to permeabilization with wash/permeabilization buffer (0.05% saponin (Sigma-Aldrich; Cat# SAE0073) diluted in PBS) and staining for additional immunocytochemical endpoints. Following immunostaining for other proteins of interest, cells were imaged using a PerkinElmer Opera Phenix high content microscope at 20X magnification. Number of EdU-positive nuclei was quantified over 25 FOVs per condition per biological replicate.

### pSIVA survival assay

Annexin-based pSIVA(TM) (Biotechne; Cat# NBP2-29382) probes were used to quantify cell survival. Briefly, fresh BMDM medium containing pSIVA (1:100) and NucBlue Live ReadyProbes (Invitrogen; Cat# R37605) without supplemental M-CSF was added to cells 24 hours after plating immediately prior to the addition of various lysosomal drugs. After 30 minutes, lysosomal drugs were spiked in and cells were imaged using a PerkinElmer Opera Phenix high content microscope at 63X magnification. Average # of pSIVA-positive cells/well was quantified over 23 FOVs per condition per biological replicate.

### Zymosan uptake assay of phagocytosis

Phagocytosis was assessed by measuring uptake of fluorescent zymosan particles. For pharmacological experiments, cells were pre-treated with lysosomal drugs for 22 hours. On the next day, Zymosan A Bioparticles conjugated to Alexa Fluor 488 (Invitrogen; Cat# Z23373) were reconstituted at 2mg/mL in BMDM medium and probe sonicated. Fresh BMDM medium (without supplemental M-CSF) containing NucBlue Live ReadyProbes and 50μg/mL Zymosan-488 was added to each well. Lysosomal drugs were immediately re-added, and plates were incubated at 37°C and 5% CO_2_. After 6 hours, cells were washed twice with PBS to remove excess extracellular zymosan and imaged in Live Cell Imaging Solution (Invitrogen; Cat# A14291DJ) using a PerkinElmer Opera Phenix high content microscope at 63X magnification. Mean zymosan-488 spot area per cell was quantified over 55 FOVs per condition per biological replicate.

### Transwell chemotaxis migration assay

For transwell migration assays, cells were plated onto Cultrex poly-L-lysine (R&D Systems; Cat# 3438-100-01)-coated 6-well plates at a density of 1.25 million cells per well in fresh BMDM media (with 50ng/mL M-CSF supplementation). After 24 hours, fresh BMDM media containing lysosomal drugs was added and cells were treated for 22 hours. Then, treated BMDMs were lifted using pre-warmed Detachin (Genlantis; Cat# T1000110), centrifuged at 400xg for 5 minutes, resuspended in Chemotaxis Assay Buffer [phenol red-free RPMI-1640 (Gibco; Cat# 32404014), 0.1% BSA] containing 5μM Calcein AM (Corning; Cat# 354217) at a density of 1 million cells/mL, and incubated at 37°C and 5% CO2 for 45 minutes. Labeled cells were then pelleted at 400xg for 5 minutes and resuspended in Chemotaxis Assay Buffer at a density of 1 million cells/mL. Next, 30,000 labeled cells from each sample were added into the top compartment of a Corning FluoroBlok 24-Multiwell insert system with an 8.0μM high density PET membrane (Corning; Cat# 351158) and slowly lowered into plate containing 700mL Chemotaxis Assay buffer with or without 100ng/mL mouse recombinant C5a (Bio-Techne; Cat# 2150-C5). Next, 485/530nm (Ex/Em) wavelength fluorescent values were read at 10 minute intervals for 80 minutes total using a Synergy Neo2 microplate Reader (BioTek). Migration index was calculated by subtracting the fluorescence values of the vehicle well from the paired well containing C5a chemoattractant, and dividing by the fluorescence from the vehicle well to normalize for any minor differences in cell loading across replicates and conditions.

### CRISPR/Cas9-mediated knockout of target genes

CRISPR/Cas9 was used to selectively knockout target genes in mouse BMDMs as previously described^66^, with modifications. Briefly, WT mouse BMDMs were thawed and plated on uncoated 6-well tissue culture-treated dishes at a density of 2 million cells per well in BMDM medium supplemented with 50ng/mL M-CSF. On the following day, ribonucleoproteins (RNPs) containing 1.5μL of 100μM single-guide RNA pools (sgRNAs) (Synthego; equimolar concentrations of 3 unique sgRNAs for each target gene per pool) and 1.5μL of 20μM SpCas9 2NLS (Synthego) per reaction were assembled. BMDMs were lifted using Detachin (Genlantis; Cat# T1000110), centrifuged at 400xg for 5 minutes, resuspended in BMDM medium containing M-CSF, and counted. Cell suspensions were separated into individual tubes containing 500,000 cells per reaction and pelleted again at 400xg for 5 minutes. Following centrifugation, the supernatant was removed, cell pellets were resuspended in 20μL P3 nucleofection buffer with supplement (Lonza; Cat# V4SP-3960), added to the RNP mix containing sgRNAs and SpCas9, and transfected using program CM-137 on the 4D-Nucleofector Platform (Lonza; Cat# AAF-1003S). Immediately after transfection, 180μL of BMDM medium containing 50ng/mL M-CSF was added to each reaction, cells were mixed by pipetting 3 times, and transfected cells were plated evenly across 10 wells of poly-D-lysine-coated 96-well PhenoPlates (PerkinElmer; Cat# 6055508). Half-media changes of BMDM medium containing 50ng/mL M-CSF were performed every other day and endpoints were assessed until 14 days after transfection.

### Statistical analyses

Statistical analyses of scRNA-seq, bulk RNA-seq, and H3K27ac ChIP-seq data were performed as described above. All other data were analyzed using Prism 10 (GraphPad). Information regarding individual statistical tests is provided in each panel’s corresponding figure legend.

## Data Availability

All raw and processed bulk and scRNA-seq data are available via the NCBI GEO repository as SuperSeries accession GSE27255. All raw and processed ChIP-seq data are available via the NCBI GEO repository as accession GSE279777.

## Code Availability

All code used to analyze the bulk RNA-seq, scRNA-seq, and ChIP-seq data is available upon request.

## ACKNOWLEDGMENTS

We thank Robert Thorne, Stacy Henry, Kate Monroe, Joe Lewcock, Mark Kafka, Erica Kratz, Christian Haass, Anja Capell, and Georg Werner for their useful comments, suggestions, and feedback. We also thank Dara Leto for helping to develop CRISPR-based approaches in BMDMs.

## AUTHOR CONTRIBUTIONS

L.T. and G.D.P. conceived and designed this study. L.T., L.L.S., C.H., M.J.S., and L.S. performed mouse scRNA-seq, and L.T., D.T., and T.S. analyzed the scRNA-seq data. L.T., L.L.S., and M.J.S. performed immunohistological staining of aged *Grn* KO mouse brains, and J.C.D. developed the pipeline for automated analysis of the tissue. C.B. and C.K.G. performed mouse H3K27ac ChIP-seq analysis. L.T., A.R., E.W.S., and C.M.H. collected primary microglia and BMDMs for *in vitro* experiments. L.T., C.H, and G.L.d.M. performed bulk RNA-seq, and L.T., D.T., and T.S. ran analyses. A.R. performed super-resolution microscopy of tissue and cells. L.T. and G.A.F. performed lysosomal pH measurements. L.T. performed all other experiments, with assistance from L.L.S., A.R., E.W.S., C.M.H., and E.L. L.T. and G.D.P. wrote the manuscript. All authors edited the manuscript and provided comments.

## ADDITIONAL INFORMATION

**Extended data** is available for this paper upon request.

**Supplementary information:** The online version contains supplementary material available upon request.

**Correspondence and requests for materials** should be addressed to and will be fulfilled by Gilbert Di Paolo.

## Extended Data Figure Legends

**Extended Data Figure 1.**
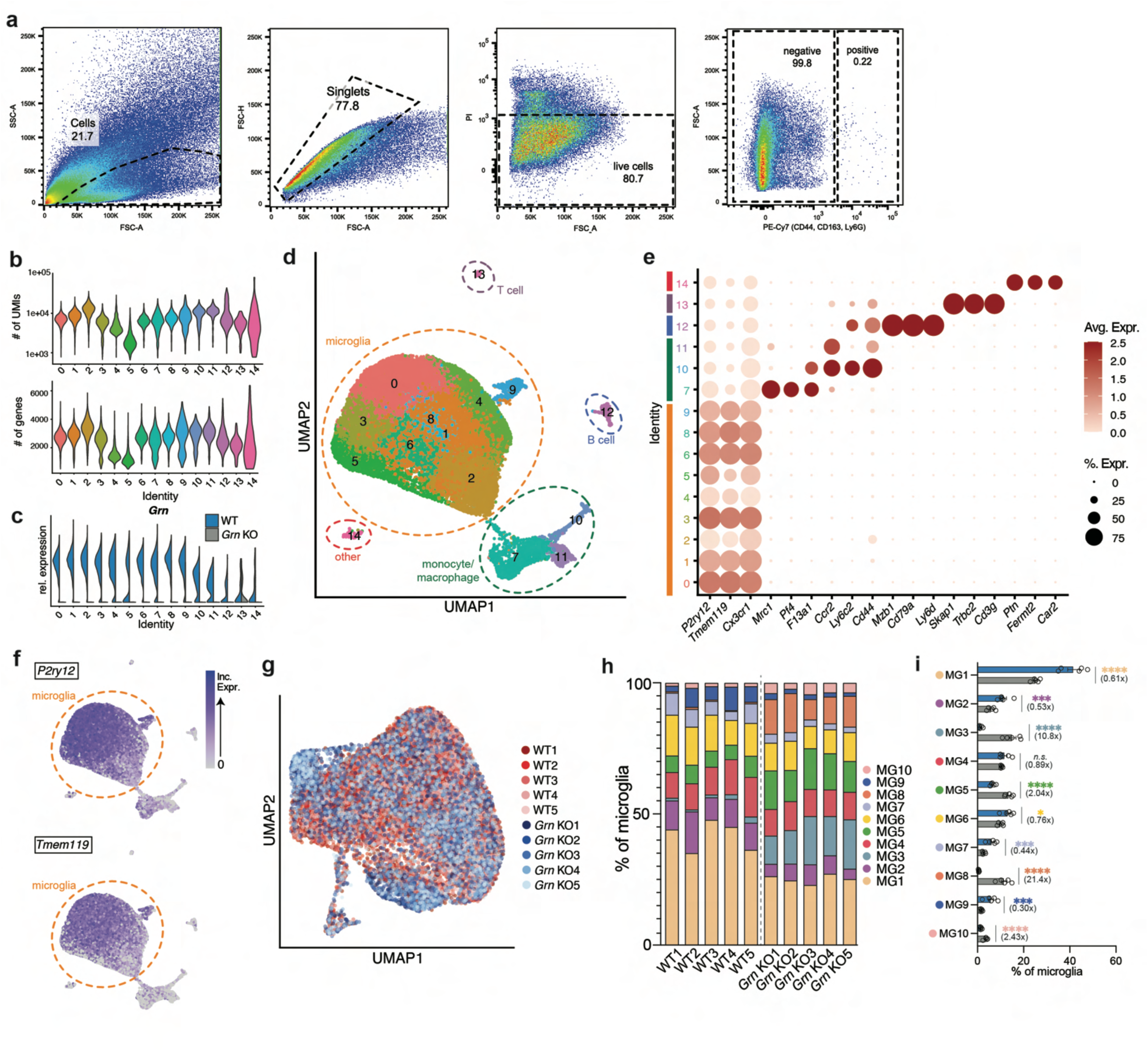
Quality control for scRNA-seq of aged *Grn* KO microglia. **(a)** Representative FACS plots displaying gating strategy for selecting live, CD44/CD163^-^/Ly6G^-^ cells for subsequent scRNA-seq. **(b)** Violin plots displaying # of unique molecular identifiers (UMIs) and # of genes detected in each cluster of the scRNA-seq dataset. **(c)** Violin plot displaying relative *Grn* transcript expression across each cluster of the scRNA-seq dataset, split by genotype. **(c)** UMAP of clusters identified among all high-quality cells of the scRNA-seq dataset. **(e)** Dot plot of marker genes for different clusters. **(f)** UMAP displaying relative expression of selected microglial markers of interest used to annotate microglial subclusters. **(g)** UMAP displaying integration of microglia from all individual mouse replicate samples. **(h)** Bar graph displaying percentages of different microglial subclusters in each sample. **(i)** Bar graphs displaying relative abundance and enrichment of different microglial subcluster (n=5 biological replicates/genotype). Data presented in (i) are mean±s.e.m. The *propeller* R package was used to compare the relative abundance of each cluster across genotypes in (i). \**P*<0.05, \*\*\**P*<0.001, \*\*\*\**P*<0.0001.

**Extended Data Figure 2.**
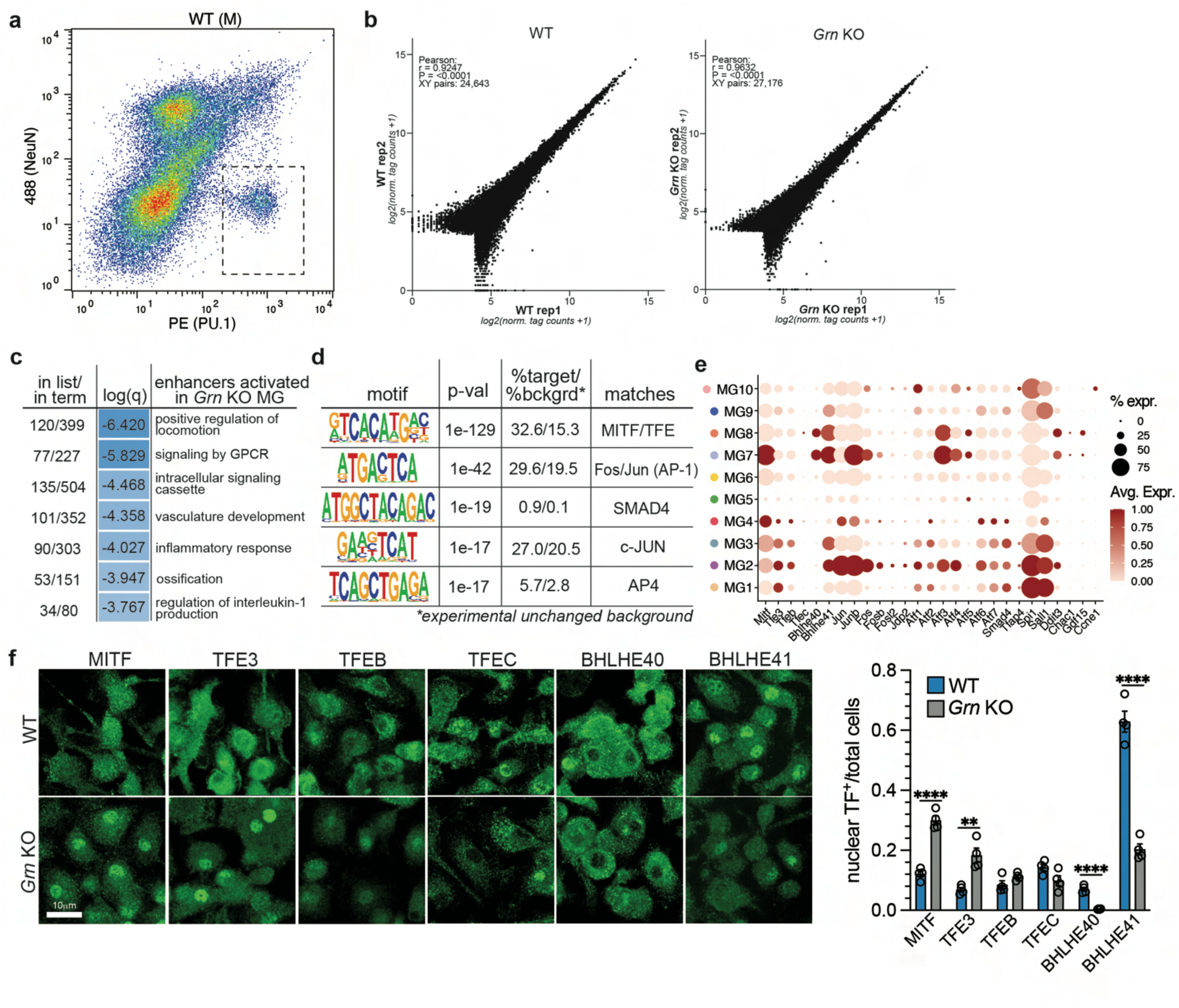
Quality control for H3K27ac ChIP-seq of aged *Grn* KO microglia and analysis of candidate transcription factors. **(a)** Representative FACS plots displaying comparable proportions of Pu.1^+^/NeuN^-^/Olig2^-^ nuclei collected for H3K27ac ChIP-seq from WT two year old mice (see Fig. 2). **(b)** Scatter plots displaying similarity of H3K27ac peaks across biological replicates for WT (left) and *Grn* KO (right) samples. **(c)** Gene ontology analysis of activated enhancers in aged *Grn* KO microglia. **(d)** Motif enrichment analysis of activated enhancers in *Grn* KO microglia relative to an experimental unchanged background. **(e)** Dot plot displaying relative expression of putative regulatory transcription factors in the different microglia subclusters from the scRNA-seq dataset. **(f)** Representative images and quantification of transcription factor localization in WT and *Grn* KO BMDMs in culture. Data presented in (f) are mean±s.e.m. Student’s t-tests were performed to compare across genotypes in (f). \*\**P*<0.01, \*\*\*\**P*<0.0001.

**Extended Data Figure 3.**
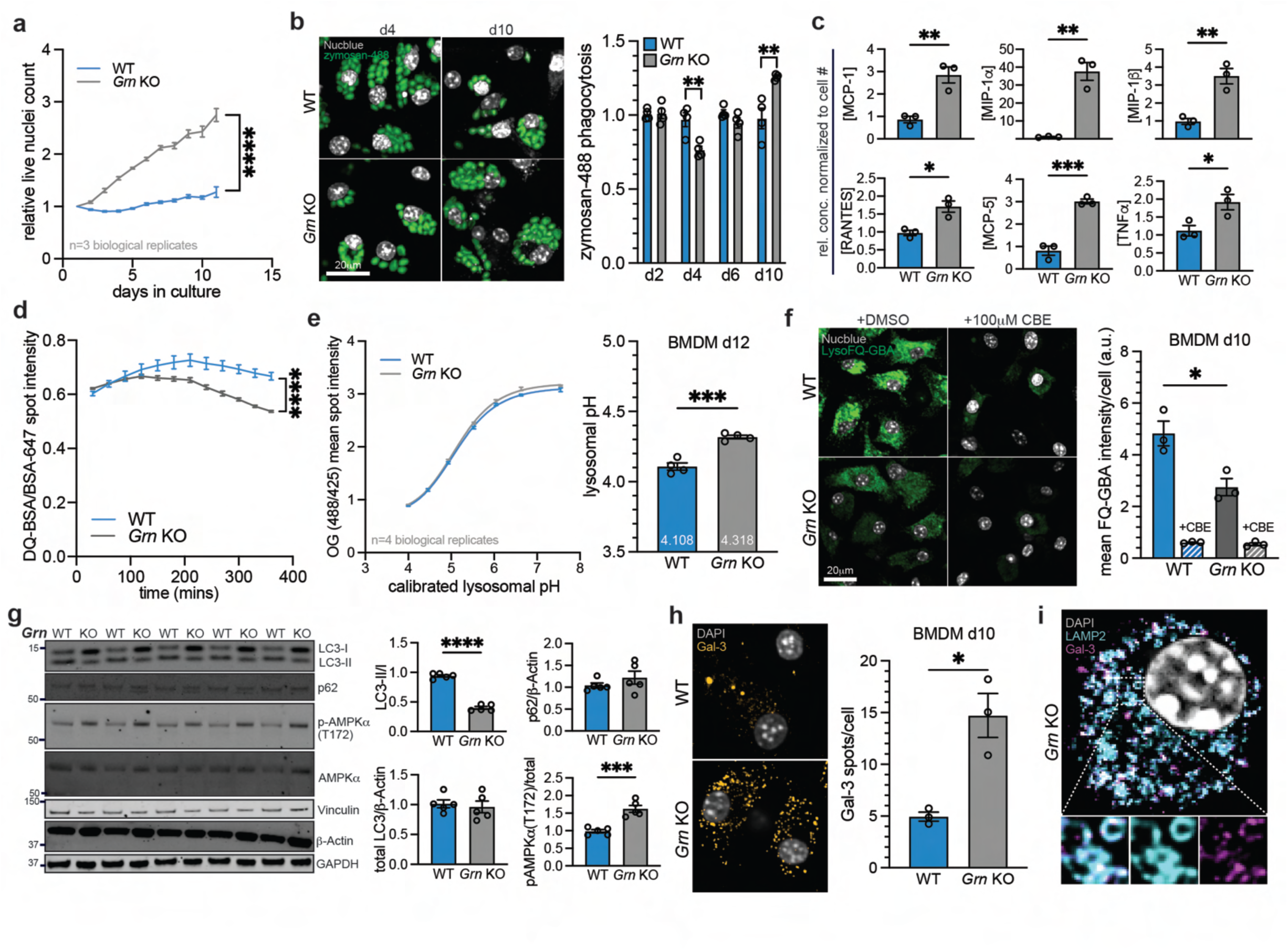
*Grn* KO BMDMs display impairments in diverse lysosomal properties and myeloid cell functions. **(a)** Quantification of live nuclei count over time in WT and *Grn* KO BMDMs after plating (n=3 biological replicates/genotype). **(b)** Representative images and quantification of zymosan-488 phagocytosis (spot area/cell normalized to WT) in WT and *Grn* KO BMDMs (n=4 biological replicates/genotype). **(c)** Measurement of secreted pro-inflammatory cytokines/chemokines by WT and *Grn* KO BMDMs (n=3 biological replicates/genotype; normalized to cell # and to WT). **(d)** Measurement of lysosomal proteolysis over time (post-loading probe) in WT and *Grn* KO BMDMs using DQ-BSA/BSA-647 (n=4 biological replicates/genotype). **(e)** Calibration curve and measurement of lysosomal pH in WT and *Grn* KO BMDMs (n=4 biological replicates/genotype). **(f)** Representative images and measurement of GCase activity in WT and *Grn* KO BMDMs with and without GCase inhibitor CBE (n=3 biological replicates/genotype). **(g)** Western blots and quantification of key autophagy-related proteins in d10 WT and *Grn* KO BMDMs (n=5 biological replicates/genotype). **(h)** Representative images and quantification of Gal-3 spots per cell in WT and *Grn* KO BMDMs to measure lysosome membrane permeabilization (n=3 biological replicates/genotype). **(i)** Representative super-resolution image of *Grn* KO BMDMs displaying co-localization between Gal-3 and lysosome marker Lamp2. Data presented in (a-e, g-i) are mean±s.e.m. Two-way *ANOVA* tests with multiple comparisons were performed to compare across genotypes in (a, d). Student’s t-tests were performed to compare across genotypes in (b,c, e-h). \**P*<0.05, \*\**P*<0.01, \*\*\**P*<0.001, \*\*\*\**P*<0.0001.

**Extended Data Figure 4.**
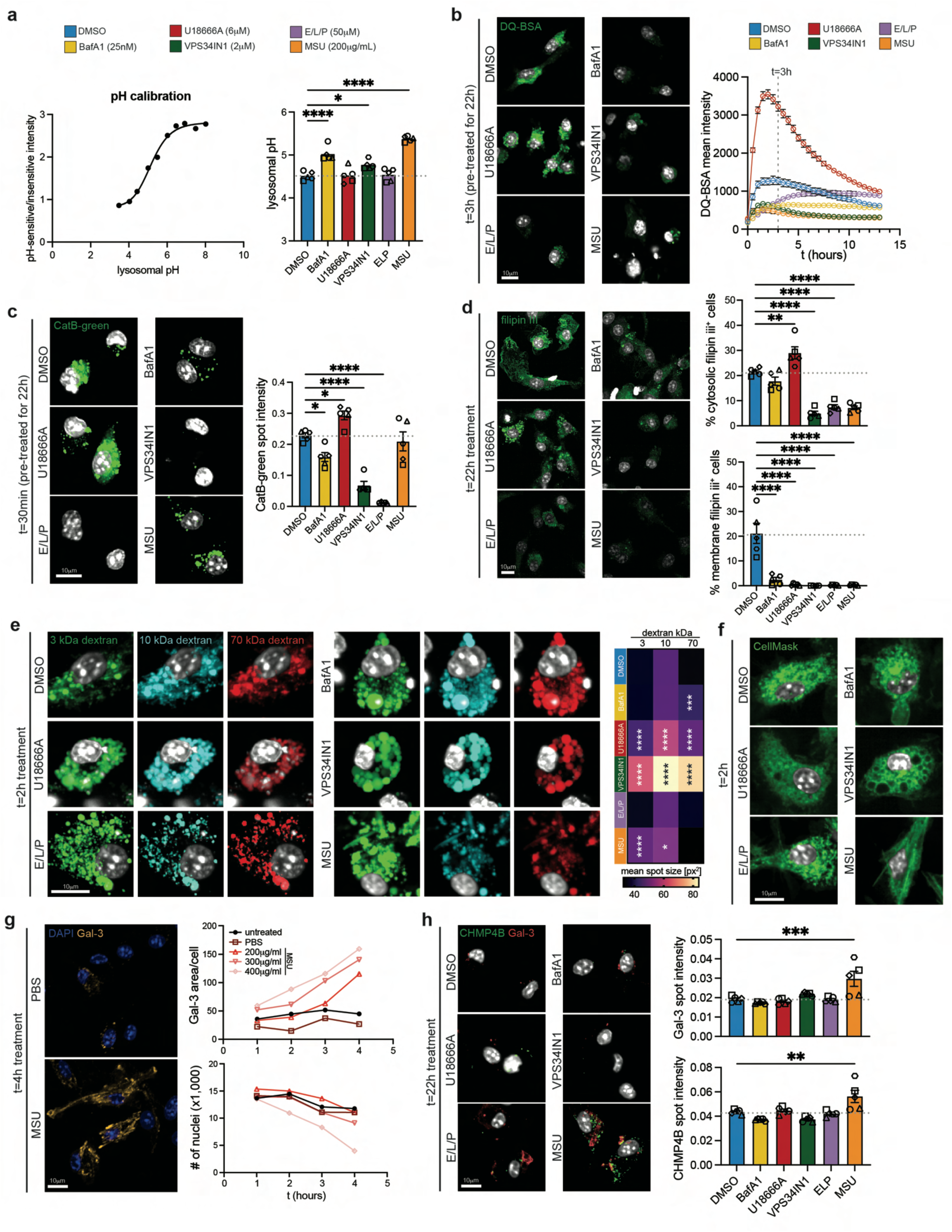
Validation of pharmacological perturbations of lysosomal properties in BMDMs. **(a)** Calibration (left) and measurement (right) of lysosomal pH in WT BMDMs treated with various lysosomal drugs for two hours (n=5 biological replicates/genotype). **(b)** Representative images (t=3h) and longitudinal measurement of lysosomal proteolysis using DQ-BSA in WT BMDMs following 22 hour treatment with various lysosomal drugs (n=5 biological replicates/genotype). **(c)** Representative images and measurement of lysosomal proteolysis using CatB-green at t=30min in WT BMDMs following 22 hour treatment with various lysosomal drugs (n=5 biological replicates/genotype). **(d)** Representative images and measurement of cholesterol using filipin iii in WT BMDMs following 22 hour treatment with various lysosomal drugs (n=5 biological replicates/genotype). **(e)** Representative images and measurement of endolysosome size in WT BMDMs loaded with different sized dextran particles, following a two hour treatment with various lysosomal drugs (n=5 biological replicates/genotype). **(f)** Representative images displaying vacuolation following treatment with certain lysosomal drugs. **(g)** Representative images and quantification of Gal-3 area per cell and nuclei count following treatment with increasing concentrations of monosodium urate (MSU) crystals for increasing durations (n=3 biological replicates/genotype) to measure lysosome membrane permeabilization (n=1 biological replicate per concentration per timepoint). **(h)** Representative images and measurement of lysosome membrane permeabilization in WT BMDMs following a 22 hour treatment with various lysosomal drugs (n=5 biological replicates/genotype). Data presented in (a-d,h) are mean±s.e.m. One-way *ANOVA* tests with multiple comparisons were performed to compare across genotypes in (a-e,h). \**P*<0.05, \*\**P*<0.01, \*\*\**P*<0.001, \*\*\*\**P*<0.0001.

**Extended Data Figure 4.**
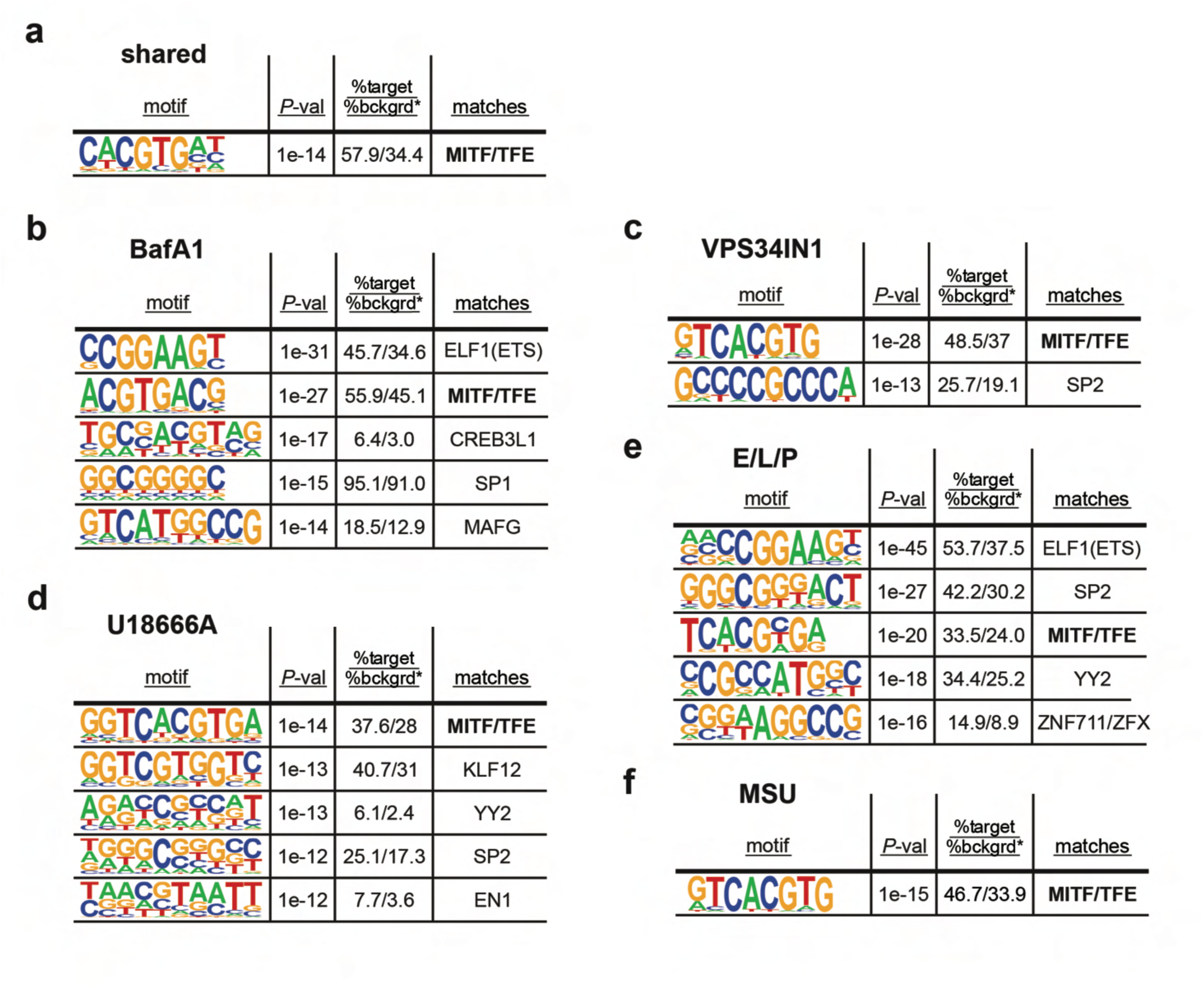
*De novo* motif enrichment analysis of promoter regions of genes upregulated in BMDMs following pharmacological perturbations of lysosomal properties. **(a)** The single significantly enriched *de novo* motif in the promoter regions (-2kb-+500bp) of the common set of 276 genes upregulated by all five pharmacological perturbations. **(b)** The top five significantly enriched *de novo* motifs in 2,859 promoter regions of genes upregulated by treatment with BafA1 (FDR <0.05). **(c)** The two significantly enriched *de novo* motifs in 2,452 promoter regions of genes upregulated by treatment with VPS34IN1 (FDR <0.05). **(d)** The top five significantly enriched *de novo* motifs in 1,459 promoter regions of genes upregulated by treatment with U18666A (FDR <0.05). **(e)** The top five significantly enriched *de novo* motifs in 2,013 promoter regions of genes upregulated by treatment with E/L/P (FDR <0.05). **(f)** The single significantly enriched *de novo* motif in the 951 promoter regions of genes upregulated by treatment with MSU crystals (FDR <0.05).

**Extended Data Figure 6.**
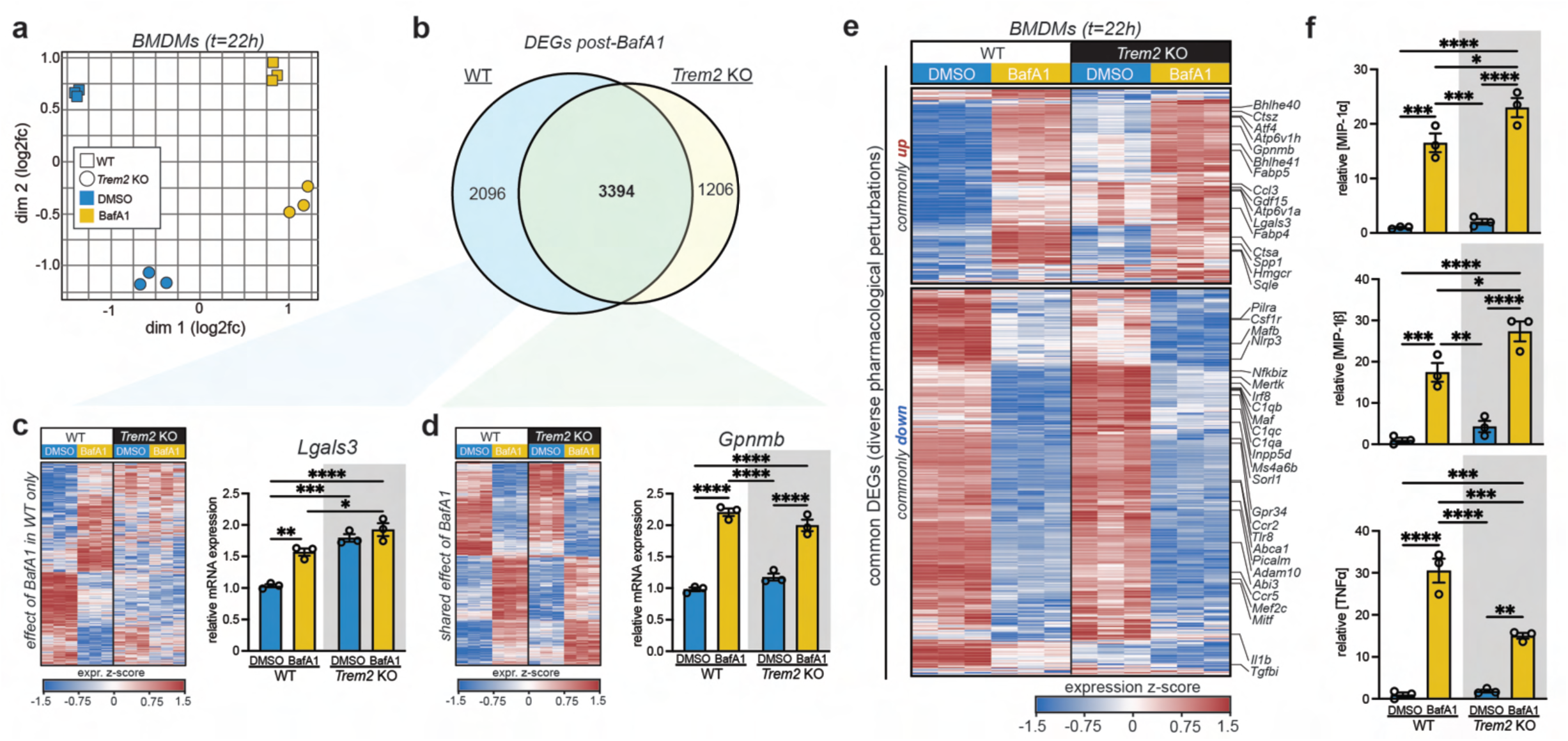
TREM2 is not require for myeloid cell response to lysosomal deacidification. **(a)** Multidimensional scaling plot displaying overall transcriptional (dis)similarity between WT and *Trem2* KO BMDMs treated with DMSO or BafA1 for 22 hours. **(b)** Venn diagram displaying overlap of DEGs in WT and *Trem2* KO BMDMs treated with BafA1 for 22 hours. **(c)** Heatmap and bar plot displaying DEGs that are observed in WT BMDMs and not *Trem2* KO BMDMs (n=3 biological replicates/condition). **(d)** Heatmap and bar plot displaying DEGs that are observed in both WT BMDMs and *Trem2* KO BMDMs (n=3 biological replicates/condition). **(e)** Heatmap displaying high conservation of common transcriptional response to lysosomal dysfunction in *Trem2* KO BMDMs (n=3 biological replicates/condition). **(f)** Secretion of key pro-inflammatory chemokines and cytokines upon BafA1 treatment is conserved in *Trem2* KO BMDMs (n=3 biological replicates). Data presented in (c,d,f) are mean±s.e.m. One-way *ANOVA* tests with multiple comparisons were performed to compare across groups in (c,d,f). \**P*<0.05, \*\**P*<0.01, \*\*\**P*<0.001, \*\*\*\**P*<0.0001.

**Extended Data Figure 7.**
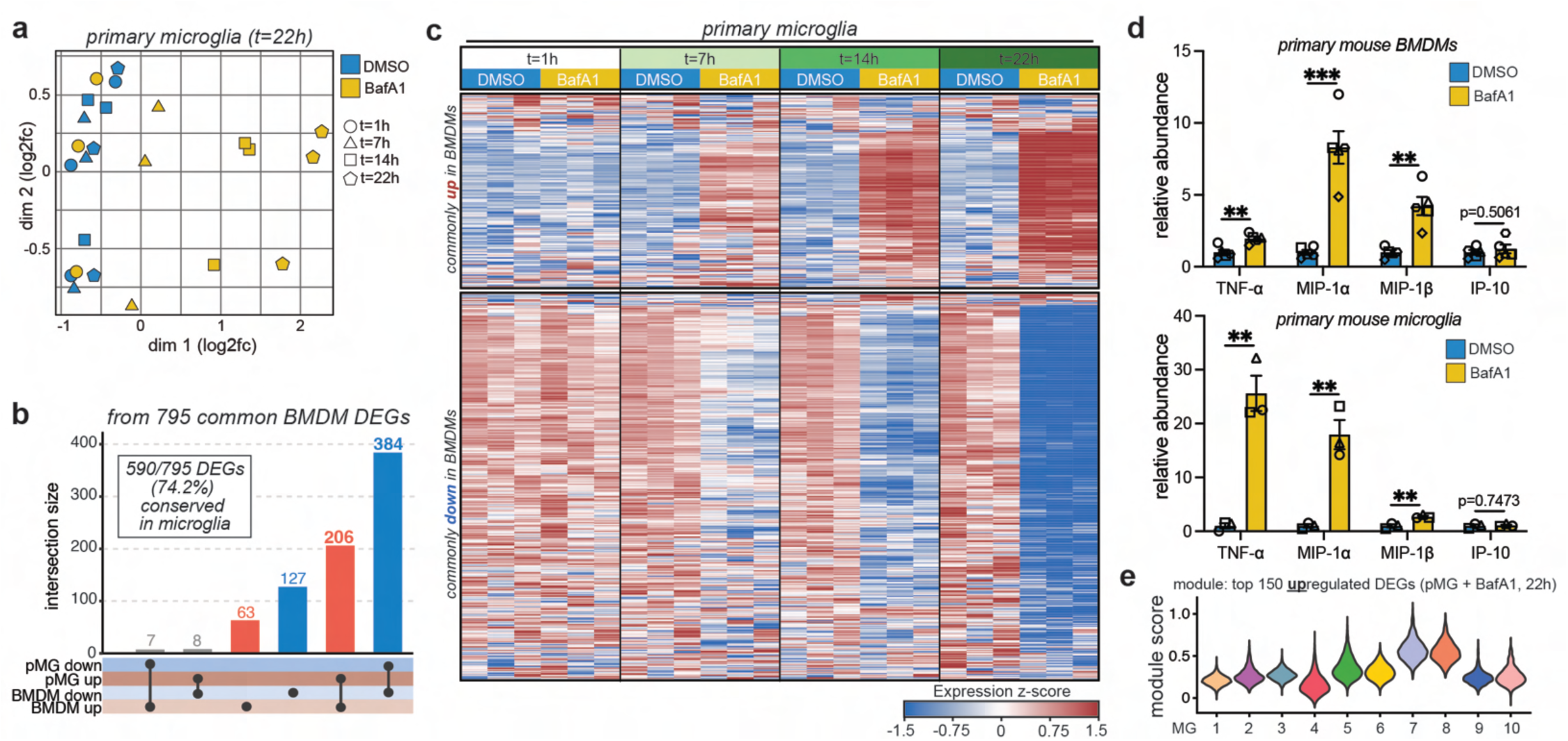
Myeloid cell response to lysosomal deacidification is highly conserved in primary microglia. **(a)** Multidimensional scaling plot displaying overall transcriptional (dis)similarity between WT primary microglia (pMG) treated with DMSO or BafA1 for 1,7,14, or 22 hours. **(b)** Bar graph displaying high degree of concordance in directionality of common DEGs between BMDMs and pMG. **(c)** Heatmap displaying high conservation of common transcriptional response to lysosomal dysfunction in pMG (n=3 biological replicates/condition). **(d)** Secretion of key pro-inflammatory chemokines and cytokines upon BafA1 treatment is conserved across BMDMs and pMG (n=3-4 biological replicates). **(e)** Average expression of top 150 upregulated genes [from WT treated with BafA1 *in vitro*] in each microglia subcluster from aged *Grn* KO microglia scRNA-seq. Data presented in (d) are mean±s.e.m. Student’s t-tests were performed to compare across conditions in (d). \*\**P*<0.01, \*\*\**P*<0.001.

**Extended Data Figure 8.**
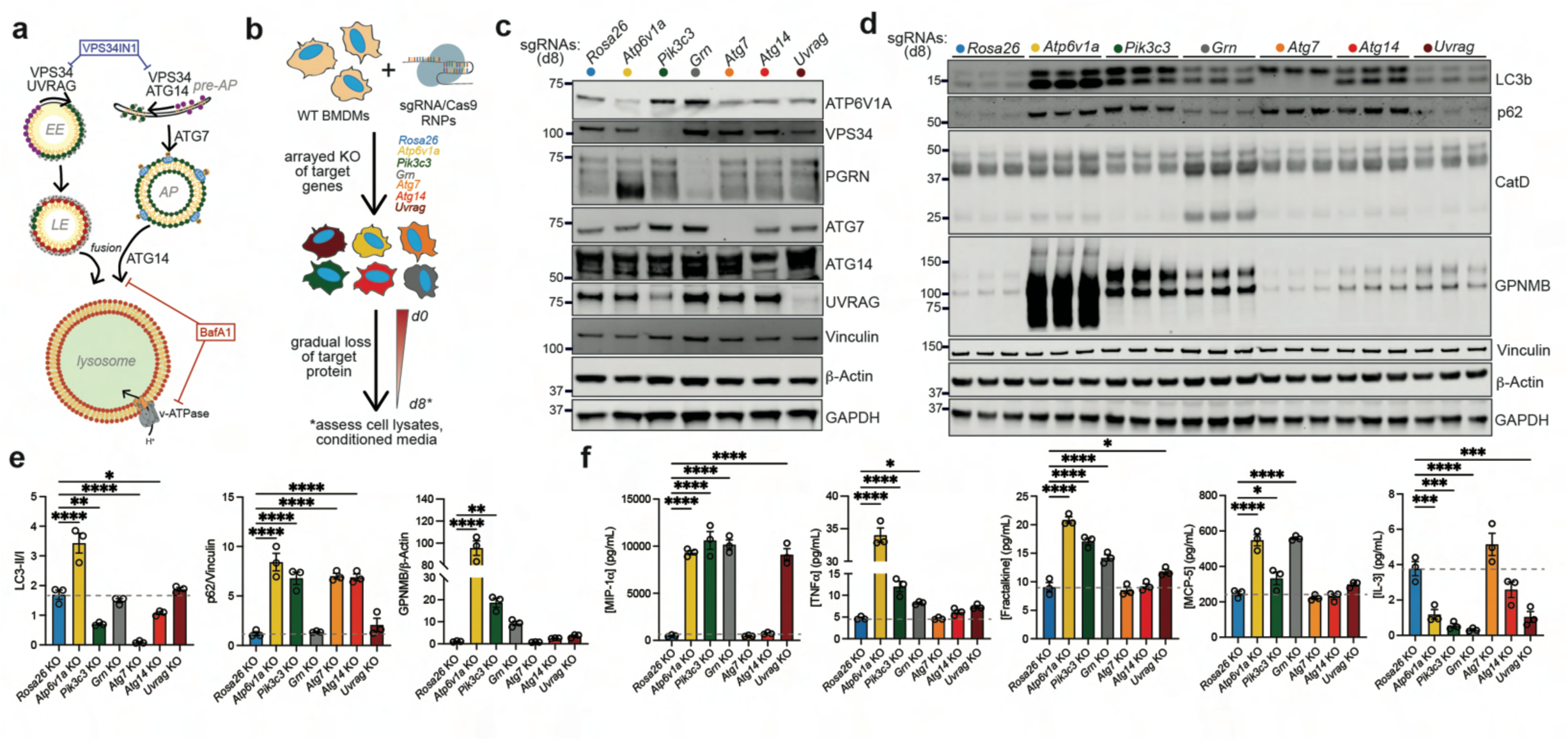
Canonical macroautophagy impairment does not phenocopy *Grn* KO BMDM phenotypes. **(a)** Schematic displaying role of various ATG/VPS proteins in endosome and autophagosome maturation/fusion. **(b)** Schematic displaying approach to genetically perturb various autophagosome formation/maturation using CRISPR/Cas9 RNP delivery in WT BMDMs. **(c)** Western blots demonstrating successful reduction of target proteins of interest by day 8 post-CRISPR. **(d,e)** Western blot and quantification confirming successful impairment at various stages during autophagosome formation and maturation (n=3 biological replicates/genotype). **(e)** Quantification of total cellular GPNMB in BMDMs from each genotype (n=3 biological replicates/genotype). **(f)** Quantification of secreted cytokines from BMDMs from each genotype (n=3 biological replicates/genotype). Data presented in (e-g) are mean±s.e.m. One-way *ANOVA* tests with multiple comparisons were performed to compare across groups in (e-g). \**P*<0.05, \*\**P*<0.01, \*\*\**P*<0.001, \*\*\*\**P*<0.0001.

## SUPPLEMENTARY TABLE LEGENDS

Supplementary Table 1. scRNA-seq of microglia from aged *Grn* KO mice.

Sample metadata and differential expression to identify marker genes between different microglia subclusters.

Supplementary Table 2. H3K27ac ChIP-seq of microglia from aged *Grn* KO mice.

Sample metadata and analysis of differential enhancer activity between WT and *Grn* KO microglia.

Supplementary Table 3. Bulk RNA-seq of constitutive *Grn* KO pMG and BMDMs.

DEG lists between WT and *Grn* KO pMG or BMDMs. Genes with FDR < 0.1 and absolute fold change > 20% were considered significantly differentially expressed.

Supplementary Table 4. Bulk RNA-seq of WT BMDMs treated with various lysosomal drugs.

DEG lists between DMSO-treated and drug-treated WT BMDMs over time. Genes with FDR < 0.1 and absolute fold change > 20% were considered significantly differentially expressed. Common and unique gene lists for each treatment were provided.

Supplementary Table 5. Bulk RNA-seq of WT and *Trem2* KO BMDMs treated with BafA1. DEG lists between DMSO-treated and BafA1-treated WT and *Trem2* KO BMDMs. Genes with FDR < 0.1 and absolute fold change > 20% were significantly differentially expressed.

Supplementary Table 6. Bulk RNA-seq of WT pMG treated with BafA1.

DEG lists between DMSO-treated and BafA1-treated WT pMG. Genes with FDR < 0.1 and absolute fold change > 20% were considered significantly differentially expressed.

Supplementary Table 7. Bulk RNA-seq of WT BMDMs following CRISPR of various genes of interest.

Single-guide RNA (sgRNA) sequences used to knock out target genes. DEG lists between target gene KO and *Rosa26* KO BMDMs at day 4 and day 13 post-CRISPR. Genes with FDR < 0.1 and absolute fold change > 20% were considered significantly differentially expressed.

Supplementary Table 8. Bulk RNA-seq of WT pMG treated with rimeporide.

DEG lists between DMSO-treated and rimeporide treated WT pMG. Genes with FDR < 0.1 and absolute fold change > 20% were considered significantly differentially expressed.

